# An unconventional HxD motif orchestrates coatomer-dependent coronavirus morphogenesis

**DOI:** 10.1101/2025.10.16.682669

**Authors:** Surovi Mohona, Anil K. Shakya, Suruchi Singh, Fiona L. Kearns, Kezia Jemison, Satchal Erramilli, Debajit Dey, Enya Qing, Benjamin C. Jennings, Balraj Doray, Anthony A. Kossiakoff, Rommie E. Amaro, Thomas Klose, Tom Gallagher, S. Saif Hasan

## Abstract

Assembly of infectious coronaviruses requires spike (S) protein trafficking by host coatomer, typically via a dibasic signal in the S cytoplasmic tail. However, the human embecoviruses HKU1 and OC43, as well as the model virus MHV, lack this motif. Here we identify a conserved His-x-Asp (HxD) sequence that functions as an unconventional coatomer-binding signal. Structural and biochemical analyses show that the MHV HxD motif engages coatomer subunits through distinct conformations, while cellular imaging demonstrates its role in directing S to assembly sites with the viral M-protein. Disruption of HxD–coatomer interactions impairs S incorporation and provokes compensatory viral adaptations, including emergence of a canonical dibasic motif or mutations in M-protein. Electron microscopy further reveals profound alterations in virion surface architecture. These findings uncover HxD as a previously unrecognized coatomer-targeting motif, highlighting an unexpected flexibility in coronavirus assembly pathways and broadening understanding of the cellular machinery that shapes coronavirus morphogenesis.

## INTRODUCTION

Protein secretion allows eukaryotic cells to shape their environment and communicate with one another. This process relies on the secretory pathway, which links organelles such as the endoplasmic reticulum (ER), the ER–Golgi intermediate compartment (ERGIC), and the Golgi network through vesicular transport^1,2^. Along this route, proteins are sequentially modified and sorted, expanding proteome diversity and achieving functional partitioning^3,4^. Disruption of these tightly coordinated events compromises cellular homeostasis and is associated with diseases including autoimmunity, cancer, and developmental disorders^5–7^.

Membrane-enveloped coronaviruses exploit the diversity in this secretory pathway to assemble infectious particles. The virion consists of three primary structural proteins^8–15^: spike (S), evident as a “corona” of outward projections from virion membranes; nucleocapsid (N), which forms intravirion ribonucleoprotein (RNP) complexes with genomic RNA, and membrane (M), which generates the framework of the virion membrane and also associates with S and RNPs to organize virus assembly **(Fig. 1A)**. Assembly takes place on intracellular ERGIC membranes^11^. A fourth minor envelope (E) protein facilitates assembled virus budding into the ERGIC and egress of the intralumenal viruses via unconventional secretory processes^16^.

**Figure 1.**
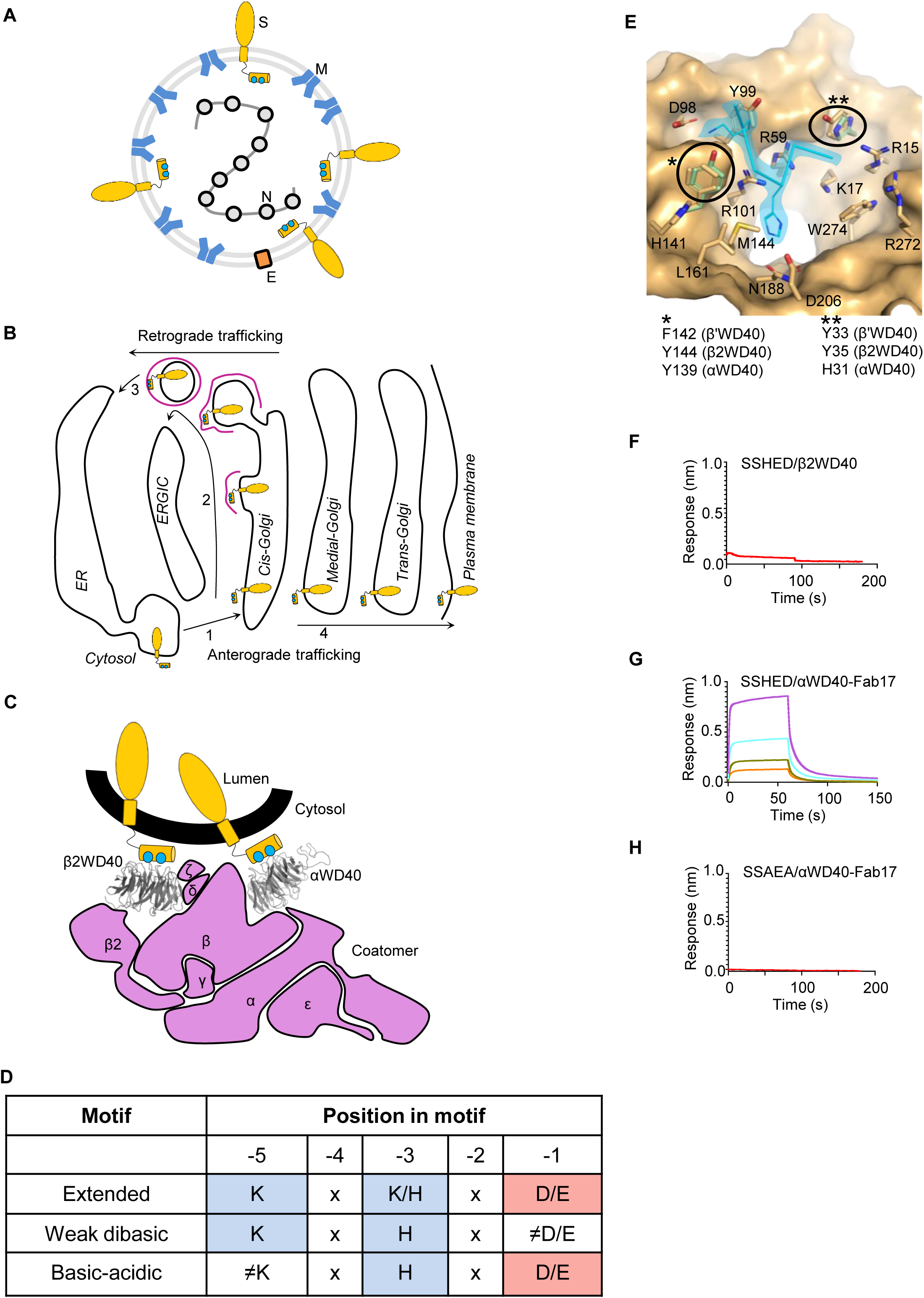
HxD motif in the MHV S tail mediates selective binding to αWD40. **(A)** Schematic of coronavirus particle showing structural proteins- S, M, N, E. For simplicity, the S molecule is shown as a monomer and not a trimer. **(B)** Schematic of S protein trafficking (yellow) with coatomer shown in magenta. **(C)** Engagement of the dibasic motif (cyan) in the S tail by coatomer αWD40 and β2WD40 domains. **(D)** Cytosolic tail motifs in endogenous coatomer clients and coronavirus S proteins. Blue, basic residue; red, acidic residue. **(E)** Binding site residues in αWD40, yeast β′WD40, and murine β2WD40, with a reference KxH peptide shown in cyan (from PDB ID 8ENW). **(F)** BLI showing no detectable interaction between the 21-mer SSHED peptide and β2WD40 (5 µM). **(G)** BLI demonstrating binding of the 21-mer SSHED peptide to αWD40 across indicated protein concentrations (5, 2.5, 1.25, 0.675 µM). **(H)** BLI showing no detectable interaction between the 22-mer SSAEA peptide and αWD40 (5 µM).

Although S is not required for particle formation, its incorporation is essential for virus binding to host cell receptors and fusion with host cell membranes^17–20^. S, a homotrimeric type I membrane glycoprotein with multidomain organization, is embedded in virions by a single-pass transmembrane segment and a C-terminal intravirion or cytosolic tail containing trafficking signals^21–27^. Synthesized in the ER, S acquires N-linked glycans on its ecto-domains, which are critical for trimerization and ectodomain folding^8^. During passage through the secretory pathway, N-glycan modification and proteolytic processing facilitates S conformations directing the receptor binding and membrane fusion events required for virus entry^20,28^. Hence, proper secretory biogenesis of the S protein is essential for infectious virus entry.

S must reach the ERGIC to be incorporated into virions. Prior investigations on coronaviruses such as SARS-CoV-2 have established that retrograde trafficking from the Golgi plays a major role in supplying S to the virus assembly site^25,29,30^ **(Fig. 1B)**. Many β-coronaviruses, including SARS-CoV-2, achieve this by mimicking host ER retrieval signals. Specifically, the cytosolic tail of S contains a dibasic Lys–x–His motif that resembles the Lys–x–Lys motif of ER-resident proteins^25^. These motifs are recognized by the coatomer complex, a conserved 560 kDa assembly of seven gene products–α, β, β′ (or β2), γ, δ, ε, ζ–that mediates retrograde transport from the Golgi to the ER to maintain protein homeostasis^31–33^ **(Fig. 1C, D)**. Coatomer binds dibasic motifs in the cytosolic tail of “client” membrane proteins through the N-terminal WD40 domains of its α and β′ subunits^34^ (αWD40 and β′WD40, respectively; **Fig. 1C**). These two WD40 domains are structural homologs: despite limited overall sequence similarity, they share extensive structural conservation, and their binding sites differ in only one or two residues^35,36^ (**Fig. 1E)**. Both engage the side chains of the dibasic residues in cargo proteins through conserved acidic patches^36^. In endogenous proteins, an acidic residue at the C-terminal position (Asp/Glu) often forms an “extended motif”^37^ (**Fig. 1D**). S proteins in coronaviruses such as SARS-CoV-2 diverge from this canonical extended motif by omitting the C-terminal acidic residue^29^ **(Fig. 1D)**, achieving partial mimicry sufficient to engage coatomer in *in vitro* assays but presumably allowing release at the ERGIC. Structure–function studies have shown that introducing a C-terminal acidic residue into SARS-CoV-2 S (e.g., Thr1273Glu) dramatically increases coatomer binding, inhibits S release, and prevents incorporation into assembling virus-like particles (VLPs)^29^.

Embecoviruses, which include the human coronaviruses HKU1 and OC43—typically associated with the common cold but capable of causing serious disease in immunocompromised individuals^38–40^—as well as the murine hepatitis virus (MHV) model, present a distinct case in molecular mimicry. They assemble in the ERGIC but lack the canonical dibasic motif found in other β-coronaviruses^11,37,41^. Alternative basic–acidic signatures (His–x–Asp or HxD) in their S tails mimic one of the two residues in the dibasic motif and the C-terminal acidic residue^37^ **(Fig. 1D)**, but whether these function as coatomer-recognition elements remains unknown. Moreover, the broader question of how coatomer hijacking influences coronavirus morphology and S protein organization in virions has not been addressed.

Here, we define the structural and functional basis of an unconventional HxD motif in the MHV S tail as a selective and indispensable coatomer-binding determinant. Using structural, biophysical, computational, and cell biological approaches, we show how this motif engages the αWD40 domain, primes coatomer interactions, and governs coronavirus morphogenesis and infectivity. These findings establish a unique mimicry strategy exploited by embecoviruses, expand the repertoire of coatomer clients, and provide a conceptual framework for understanding how coronaviruses co-opt host trafficking machinery for assembly.

## RESULTS

### HxD motif in the MHV S tail mediates selective binding to αWD40

We first tested whether the MHV HxD motif contributes to coatomer recognition. Biolayer interferometry (BLI) assays were used to assess direct binding of biotinylated MHV S tail peptides to purified N-terminal WD40 domains from individual coatomer subunits. The murine β2WD40 domain, differing in its binding site from yeast β′WD40 by two residues and its ortholog, αWD40, by a single residue, was recombinantly expressed in Expi293 cells and purified by affinity chromatography and size-exclusion chromatography **(Fig.1E; SI Fig. 1A; SI Table T1)** We observed negligible binding to immobilized MHV S tail 21-mer peptide, indicating a lack of interaction with β2WD40 **(Fig. 1F; SI Tables T2, T3)**.

Expression and purification of αWD40 for quantitative binding studies have historically been hindered by low yields, aggregation, and nonspecific BLI biosensor binding^35,36,42^. To overcome these challenges, we developed a stabilization strategy for the yeast αWD40 ortholog, which shares an identical peptide-binding site with mammalian αWD40. First, we supplemented purification buffers with n-dodecyl-β-D-maltopyranoside (DDM) to shield a ∼20-residue hydrophobic surface loop **(SI Fig. S1B)**. This dramatically reduced aggregation during proteolytic removal of the affinity tag by an equimolar amount of ULP1 protease at 4°C for 1-1.5 hours. Second, we generated a synthetic antibody fragment (Fab) against αWD40 by phage display using a highly diverse synthetic antibody library. This candidate, Fab17, formed a complex with αWD40 and markedly increased its solubility, enabling purification to ≥10 mg/mL without precipitation compared to 3–4 mg/mL for αWD40 alone **(SI Fig. S1C)**.

BLI assays using this stabilized αWD40–Fab17 complex confirmed robust binding to the wild-type MHV S tail peptide **(Fig. 1G**; K_D_= 0.50 ± 0.07 µM; **SI Table T3)**. Substitution of the HxD motif with AxA completely abolished binding, demonstrating that HxD is necessary for αWD40 engagement **(Fig. 1H)**. In contrast, enhancing mimicry of the extended motif by a Ser1320Lys substitution in a KSHED 21-mer peptide substantially strengthens binding interactions **(SI Fig. S1D)**.

These results establish the HxD motif as a selective molecular determinant that mediates direct recognition of the αWD40 subunit, defining a minimal coatomer-binding signature within the MHV S tail.

### HxD motif engages distinct binding environments in αWD40 and β2WD40 domains

To define the structural basis of S tail recognition by αWD40, we co-crystallized the wild-type MHV tail heptapeptide (Asn1318-Asp1324) with purified αWD40. Although multiple apo αWD40 structures have been reported^29,36,37,42^, only two co-crystal structures with cytosolic tail peptides are available—those of the endogenous client Emp47p and adenovirus E19^36^. In contrast, coronavirus S tails have been crystallized only with the weakly binding β′WD40 domain, not αWD40, likely due to high local concentrations that drive interactions in the crystallization droplets^29,36,43^. This paucity of αWD40–peptide structures likely reflects a conserved Lys residue that occupies the canonical −5 Lys binding pocket in the crystal lattice, hindering peptide access^42^.

Extensive screening with the DMSO-solubilized MHV heptapeptide yielded a 1.7 Å resolution αWD40– peptide co-crystal structure with one protein copy per asymmetric unit **(Fig. 2A; SI Table T4)**. Although the widely used yeast β′WD40 domain binds weakly in solution, it frequently crystallizes with endogenous coatomer clients or viral tails, likely due to high peptide concentration during crystallization^29,35,36,43^. Hence, to enable direct comparison, we also determined a 1.4 Å resolution structure of the peptide bound to murine β2WD40 **(Fig. 2A; SI Table T4)**. The β2WD40 asymmetric unit contained two protein molecules, but interpretable peptide density was present only in one; analyses therefore focused on this copy **(Fig. 2B)**.

**Figure 2.**
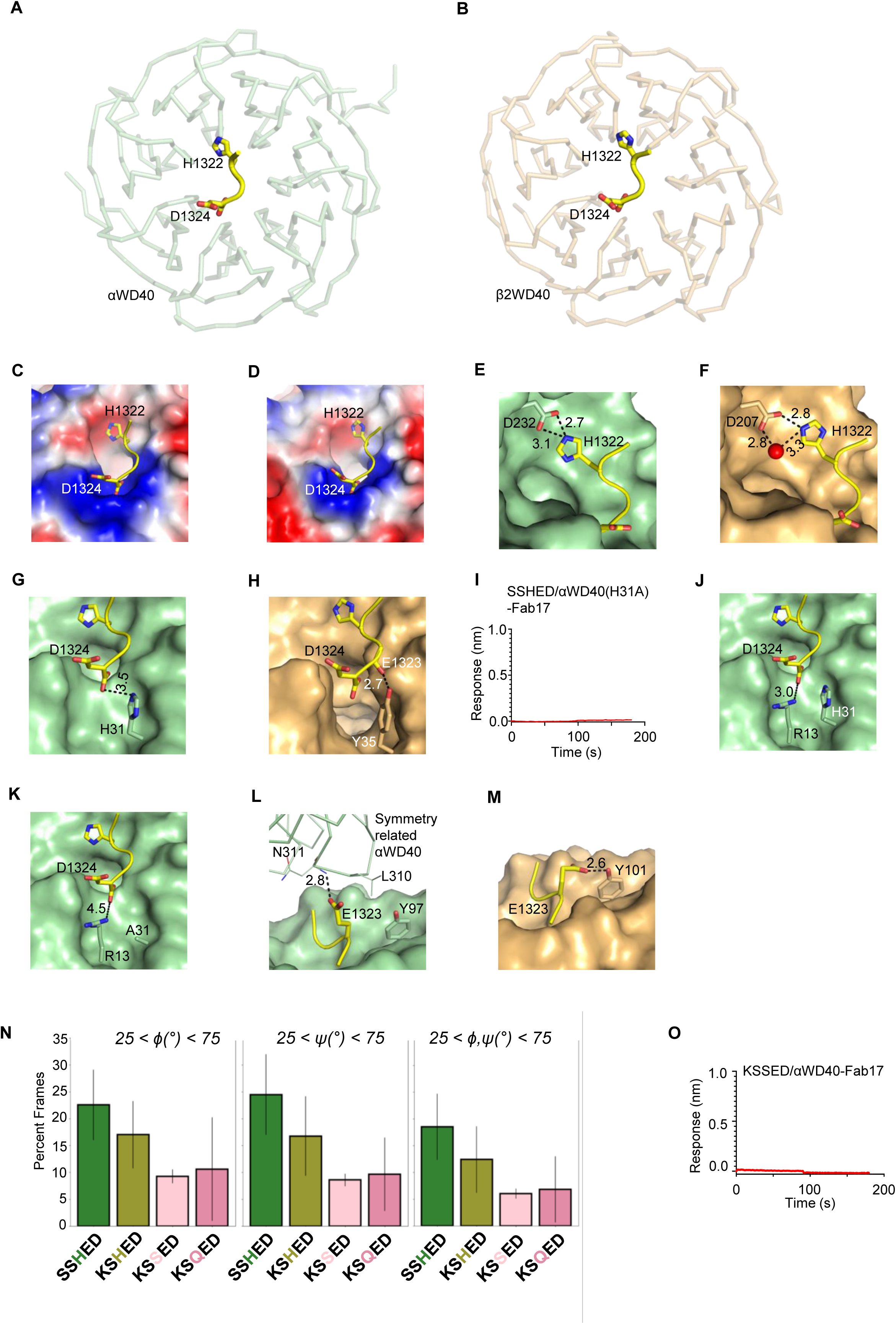
HxD motif engages distinct binding environments in αWD40 and β2WD40 domains. **(A,B)** Co-crystal structures of MHV tail peptide (Ser1321–Asp1324, highlighted) bound to αWD40 **(A)** and β2WD40 **(B)**. **(C,D)** Electrostatic surface representations of peptide-binding sites in αWD40 **(C)** and β2WD40 **(D)**; acidic residues, red; basic, blue; neutral, white. **(E,F)** Local environments of His1322 in αWD40 **(E)** and β2WD40 **(F)**; water, red sphere. **(G,H)** Substitution of αWD40 His31 with Tyr35 in β2WD40. **(I)** BLI showing loss of peptide binding upon His31Ala substitution in αWD40 (protein concentration, 5 µM). **(J,K)** Arg13 displacement in αWD40 His31Ala crystal structure **(K)** relative to wild-type αWD40 (**J;** peptide superposed from co-crystal structure in panel **K**). **(L,M)** Rotameric variation of Glu1323 side chain in αWD40 (**L**; lattice contacts) and β2WD40 (**M**; interaction with Tyr101). **(N)** Percentage of simulation frames in which the –3 residue adopted a Φ or Ψ torsion between 25° and 75°, reflecting conformational readiness for binding. **(O)** BLI showing absence of binding of KSSED peptide to αWD40 (protein concentration, 5 µM).

Both structures show the MHV heptapeptide engaging a conserved surface on αWD40 and β2WD40. Asp1324 interacts with a conserved basic cluster, and His1322 is oriented toward an acidic patch, underscoring electrostatic complementarity **(Fig. 2C, D)**. However, despite a shared overall binding mode, backbone torsions differ between αWD40- and β2WD40-bound heptapeptides, driven by four structural features. First, tail His1322 forms a direct contact with αWD40 Asp232 **(Fig. 2E)** whereas in β2WD40, a crystallographic water displaces His1322 **(Fig. 2F; SI Movie M1)**. Second, the heptapeptide-binding site differs at one position: His31 in αWD40 is replaced by Tyr35 in β2WD40, altering local electrostatics **(Fig. 2G, H)**. Mutation of αWD40 His31 to Ala abolished peptide binding **(Fig. 2I; SI Table T3)**. A 1.7 Å resolution crystal structure of this mutant showed a displacement of αWD40 Arg13 in the mutant by 1.5 Å from the acidic C-terminal carboxylate group of the heptapeptide, thus establishing the essential role of His31 in maintaining the binding site architecture **(Fig. 2J, K; SI Table T4)**. Third, Glu1323 adopts distinct rotamers: in αWD40, lattice contacts with Leu310 and Asn311 from a symmetry-related molecule engage this side-chain whereas the Glu1323 side-chain interacts with Tyr101 in β2WD40 **(Fig. 2L, M)**. Collectively, these findings elucidate the structural basis of MHV S heptapeptide interactions with αWD40 and β2WD40 domains and indicate that sequence variation and local interactions shape the conformation of the MHV heptapeptide in these complexes.

Binding of HxD and KxHxx motifs to the αWD40 domain suggests that coatomer engagement requires occupancy of only two positions within the canonical extended motif—either at –5 and –3, or –3 and –1 **(Fig. 1D)**. Whether spatial constraints between these positions further influence binding remained unclear. To address this, we performed molecular dynamics simulations with 21-mer peptides corresponding to the full-length S tail, terminating in SSHED (wild type), high-affinity KSHED, and KSSED or KSQED (−5 and −1 positions conserved, −3 replaced with polar residues).

An analysis of the Ramachandran angles revealed that KSSED and KSQED peptides displayed a markedly reduced ability to adopt backbone torsional conformations compatible with αWD40 binding **(SI Fig. S2A)**. In free simulations, His1322 at the −3 position of SSHED and KSHED occupied the target conformation (25° < Φ, Ψ < 75°) in 20% and 15% of frames, respectively, whereas KSSED and KSQED peptides achieved this state in only ∼5% of frames **(Fig. 2N; SI Fig. S2B)**. Thus, the presence of His1322 promotes conformational compatibility at −3. Structural analysis further demonstrated that His1322 engages Asp232 of αWD40 **(Fig. 2E)** in a geometry that is not supported by Ser1322, which is too short, or Gln1322 that lacks charge complementarity.

Consistent with these findings, a BLI assay confirmed that the KSSED 21-mer peptide demonstrates negligible binding to αWD40-Fab17 complex **(Fig. 2O)**. Together, these data establish that intra-motif positional constraints—particularly the presence of His at −3 —are critical determinants of coatomer recognition, by combining conformational priming with electrostatic complementarity.

#### S-coatomer affinity governs MHV infectivity and genetic adaptation

Substitutions in the SARS-CoV-2 S coatomer-binding motif alter S incorporation into virus-like particles (VLPs)^29^. To test whether this principle extends to the unconventional HxD motif of MHV S protein, we generated constructs with graded coatomer affinity based on peptide BLI data: KSHED (high), SSHED (intermediate, wild type), and SSAEA (null) **(Fig. 3A).** When expressed with MHV E, M, and N proteins, these S molecules displayed distinct incorporation phenotypes. KSHED S molecules, despite strong coatomer binding, were excluded from secreted VLPs. SSAEA S molecules lacking coatomer binding were also inefficiently incorporated, whereas SSHED S molecules supported robust assembly **(Fig. 3B).** Intracellular KSHED S molecules failed to undergo cleavage into S1/S2 subunits, consistent with coatomer-mediated retention in early secretory compartments prior to the later Golgi compartments that contain host proteases. Complete MHV VLP assembly therefore requires intermediate coatomer affinity, with both excessive and absent interactions proving detrimental.

**Figure 3.**
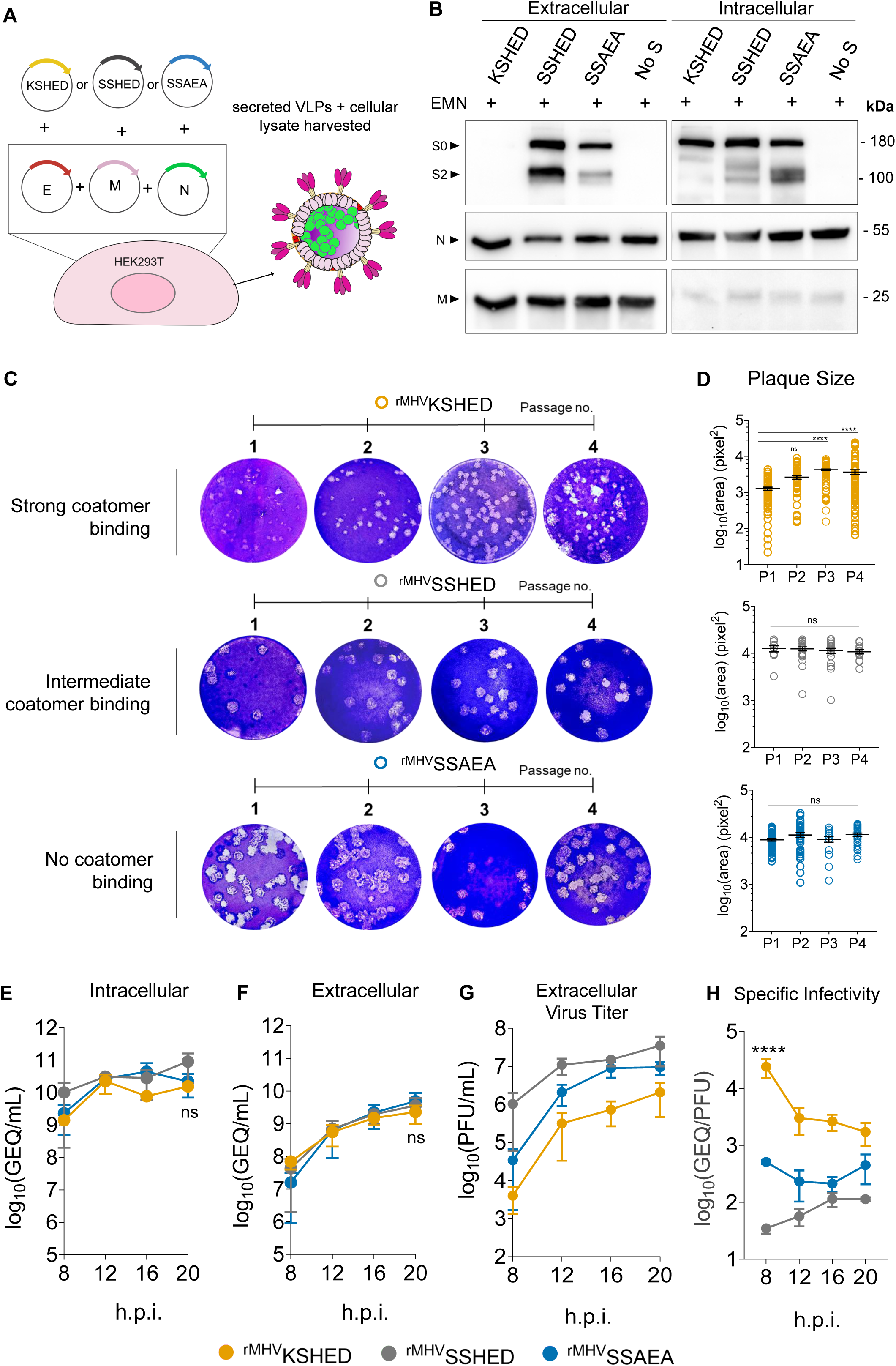
S-coatomer affinity governs MHV infectivity and genetic adaptation. **(A)** Schematic for MHV VLP assembly assay. HEK293T cells are transfected with MHV structural gene expression plasmids, including the three S plasmids-KSHED; SSHED; SSAEA. Secreted VLPs and cellular lysates harvested between 20-40 hours post-transfection. **(B)** Western blot analysis of secreted VLPs (right), and whole cell lysates (WCLs) (left) for MHV S, E, M, and N proteins. **(C)** Plaque morphology analysis of KSHED, SSHED, and SSAEA MHVs following serial passage. **(D)** Plaque area measurements performed using ImageJ. Data graphed as mean ± standard error of mean (SEM). **(E–H)** 17CL1 cells inoculated with rMHV panel at an MOI of 0.2. Extra- and intracellular samples were harvested at 8-, 12-, 16-, and 20-hours post infection. Harvested samples were analyzed via qRT-PCR for **(E)** intracellular, and **(F)** secreted genome quantitation. GEQ, genome equivalence. **(G)** Plaque assays performed using mDBT cells as indicators to quantify infectious virus titer as PFU/mL. **(H)** Specific infectivity values of secreted viruses were calculated as the ratio of GEQ to PFU. All data points are graphed as mean ± SEM and are representative of two independent experiments. Data analyzed using two-way ANOVA with Tukey’s multiple comparisons test.

Although VLPs provided valuable insights into the impact of altering affinity for the coatomer, they are limited by their inability to undergo genetic adaptation. Hence, we introduced the same coatomer-binding motif mutations into MHV S genes using a reverse genetics system based on circularized polymerase extension reaction (CPER) **(SI Fig. S3A)**. Mutations were designed to prevent same-site single-nucleotide reversions. Transfected HEK293T cells were co-cultured with MHV-permissive mDBT cells to generate recombinant (r) MHVs, which were harvested, plaque-purified, and amplified **(SI Fig. S3B).** Sequencing multiple isolates provided an assessment of virus genetic stability **(SI Table T4)**.

SSHED and SSAEA motifs remained stable during rMHV amplification. By contrast, only two of six KSHED isolates retained the engineered motif. Two acquired premature stop codons truncating C-termini of the S protein, and two carried substitutions altering coatomer-binding sequences. Interestingly, KSHED revertant viruses with substitutions (isolates c,d) and truncations (isolates e,f) displayed similar plaque morphology to SSHED and SSAEA. Infectivity analysis of isolate d (KSHET), which mimics a canonical SARS-CoV-2 KxHxT dibasic sequence with a C-terminal Thr residue, and f (Δ7C) revealed moderately attenuated infectivity, which was comparable to SSAEA **(SI Fig. S4D)**. Finally, an isolate acquired a His1322Gln substitution, thereby generating a KSQED sequence in the S tail. BLI assays supported by MD simulations above have established that such motifs do not bind the coatomer αWD40 domain. We infer that rMHVs rapidly acquired these changes to circumvent strong S–coatomer interactions.

One KSHED isolate (“KSHED a”) produced very small plaques **(Fig. 3C)** and low titers (∼10³ PFU/mL) and was highly sensitive to a single freeze–thaw cycle. Serially passaging this isolate revealed suppressor mutations that restored infectivity: plaques grew larger, titers increased, and morphology became heterogeneous **(Fig. 3C, 3D)**. By passage 4, one KSHED isolate retained the strong motif while acquiring single amino acid substitutions in NS2a and M (KSHED b; **SI Table T4**), increasing titers ∼10-fold over passage 1. Further passages yielded additional substitutions within the coatomer-binding motif (KSHED c and d; **SI Table T4**), each incrementally increasing virus titers and plaque sizes.

We next analyzed the infectivity of representative isolates that preserved the engineered motifs and carried suppressors outside S: KSHED b, SSHED a, and SSAEA a. Viral genomic RNAs and virion infectivities were quantified from m17CL1 cultures infected with each virus **(Fig. 3E-H)**. Intracellular and extracellular genomic RNAs (GEQ/mL) accumulated to similar levels across all infections **(Fig. 3E, 3F)**, indicating that S– coatomer affinity does not impair replication or virion release. In contrast, virions secreted from KSHED- and SSAEA-infected cells displayed sharply reduced infectivity (PFU/mL) relative to SSHED **(Fig. 3G)**. Specific infectivity (GEQ/PFU) analysis revealed that KSHED and SSAEA were ∼700-fold and ∼14-fold less infectious than SSHED, respectively, at 8 hours post-infection **(Fig. 3H)**. At later times post-infection, KSHED infectivity partially increased, possibly reflecting S accumulation beyond the binding capacity of coatomer complexes.

Together, these results demonstrate that mutations outside the S gene and elimination or modification of the coatomer-binding motif can partially suppress the deleterious effects of strong coatomer binding, but full restoration of infectivity requires the wild-type coatomer-binding sequence. Strong S–coatomer interactions, on the other hand, impose a potent barrier to MHV infectivity, which viruses circumvent through genetic adaptation.

#### Coatomer association controls subcellular distribution of S molecules

We examined S plasma membrane localization and the functional consequences of such localization on cell–cell fusion. Because coatomer retains S intracellularly, decreasing S–coatomer affinity was predicted to enhance syncytia formation. Consistent with this, SSAEA-virus infected cells displayed pronounced syncytia compared to SSHED-virus **(Fig. 4A, B; SI Fig. S4A-B)**. Adapted KSHED variants, including KSHET and Δ7C viruses, also induced extensive syncytia **(SI Fig. S4C)**, consistent with loss of coatomer interaction and anterograde trafficking of S to the plasma membrane for S-directed cell–cell fusion.

**Figure 4.**
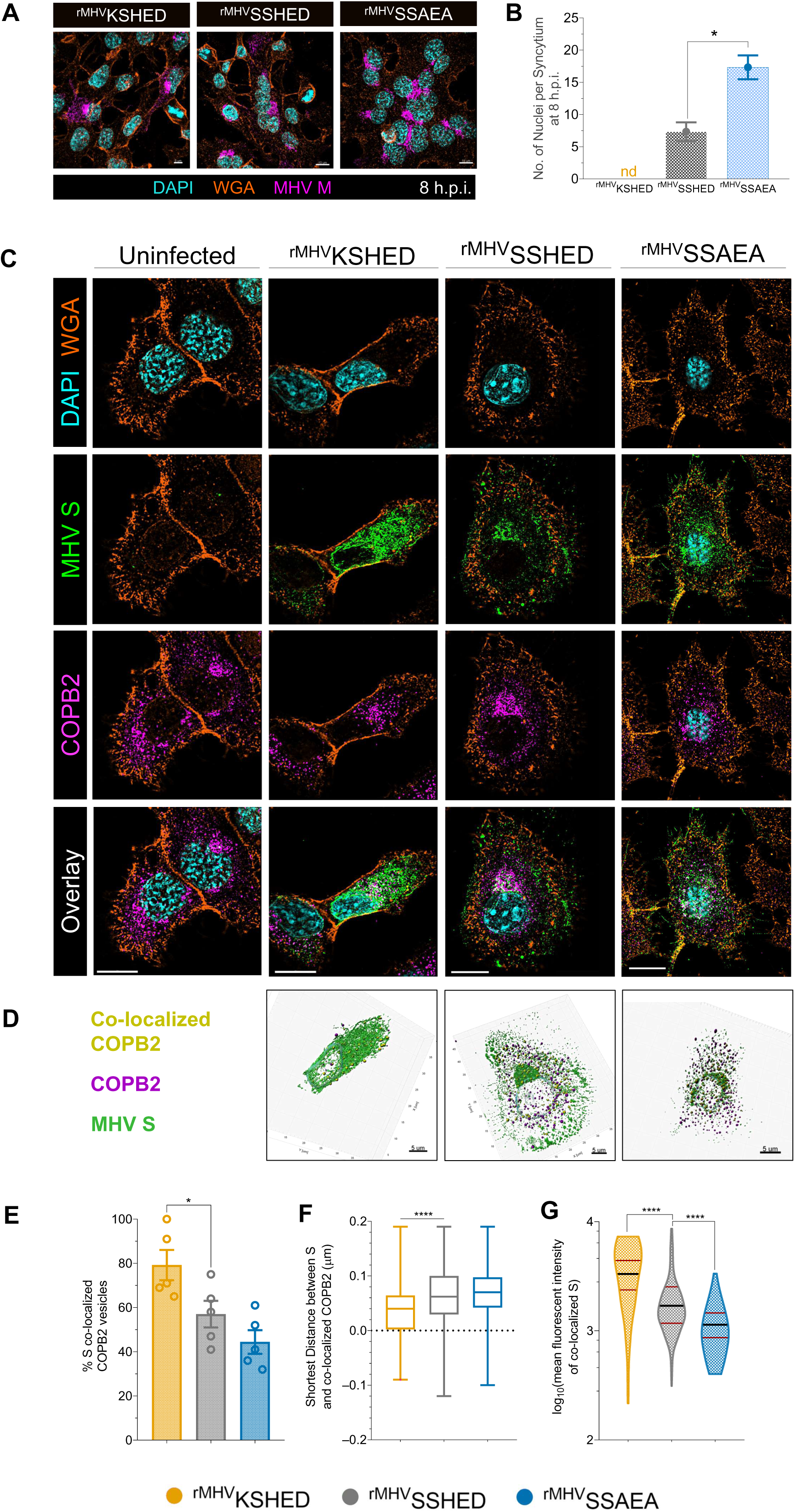
Coatomer association controls subcellular distribution of S molecules. Super-resolution immunofluorescence microscopy performed on rMHV infected cells. 17CL1 cells were infected with KSHED/ SSHED/SSAEA at an MOI of 0.5. Infected cells were fixed and permeabilized at 8-hours post infection for intracellular staining. **(A)** Images documenting cell-cell fusion (syncytia formation) at 8 hours post infection across recombinant MHV infected 17CL1 cells. **(B)** Quantitative analysis performed on 3 images per condition. *nd= no syncytia detected in KSHED infected condition at the indicated time point. Data graphed as mean ± SEM and analyzed using unpaired t-test. **(C)** MHV S stained using anti CEACAM-Fc (green), co-stained for coatomer using an antibody against COPB2 subunit (magenta). Data representative of three independent experiments. Scale bars, 10 µm. **(D)** 3D Iso-surface reconstruction of infected cells. COPB2 vesicles (magenta) within 200 nm distance of a S surface (green) were deemed co-localized (yellow). Scale bars, 5 µm. **(E)** Quantification of the percentage of COPB2 vesicles co-localized with S proteins. **(F)** shortest distance measurement between S and COPB2, and, **(G)** mean intensity of S surfaces co-localized to COPB2. Data graphed as mean ± SEM and analyzed using ordinary one-way ANOVA with Dunnett’s multiple comparisons test.

Few insights are available into how S protein impacts coatomer subcellular localization during a coronavirus infection. We addressed this question using super-resolution Lattice Structured Illumination Microscopy (Lattice SIM), which provides detailed super-resolution information on sub-cellular structures including vesicles and protein distributions^44^. Using COPB2 subunit as a marker of assembled coatomer, we observed that in uninfected, SSHED-, and SSAEA virus-infected 17CL1 cell, COPB2 puncta distributed across perinuclear and distal regions, consistent with prior immunofluorescence reports. In contrast, KSHED virus infection re-localized COPB2 vesicles to perinuclear regions **(Fig. 4C).**

S distribution mirrored these changes. S-KSHED molecules accumulated in dense perinuclear clusters, while SSHED and SSAEA S displayed both perinuclear and distal localization. Three-dimensional rendering revealed tubulovesicular S assemblies in KSHED-virus infected cells **(Fig. 4D: SI Movie M2)**. This phenotype is consistent with strong-binding S molecules trapping COPI vesicles in futile ER–ERGIC–Golgi cycling. By contrast, SSHED and SSAEA S molecules localized more broadly in infected cells, with vesicles extending toward the plasma membrane **(Fig. 4C, D; SI Movies M3, M4).**

Quantitative analysis confirmed these trends. KSHED S molecules in virus infected cells showed the highest co-localization with COPB2 vesicles (>80% vs. 40 – 60% for SSAEA and SSHED) **(Fig. 4E)**, the shortest average distances to COPB2 (40 nm vs. 50–60 nm for SSHED and SSAEA; **Fig. 4F**), and the strongest intracellular S signal intensities **(Fig. 4G)**.

These data demonstrate that the S protein requires a Goldilocks level of coatomer engagement—too strong or too weak tips the balance between retrograde and anterograde trafficking, while the right affinity ensures proper S localization for virion assembly and syncytia formation.

#### Coatomer-binding motifs dictate S–M interactions and virion composition

Coronavirus M proteins control particle assembly through their intracellular localization and interactions with S, E, and viral ribonucleoproteins^11,17^. We asked how coatomer-binding motif alterations affect intracellular S–M interactions. In SSHED-virus infected cells, M puncta predominantly co-localized with S vesicles, consistent with efficient S–M engagement. By contrast, co-localization decreased markedly in KSHED-virus infected cells (40% vs 70% in SSHED) and was modestly reduced in SSAEA-virus infections (55% S-M colocalization) **(Fig. 5A-C).** Striking differences in M distribution were also observed; M vesicles were almost exclusively distal in KSHED-virus infected cells, as opposed to both perinuclear and distal in SSHED- and SSAEA infection conditions.

**Figure 5.**
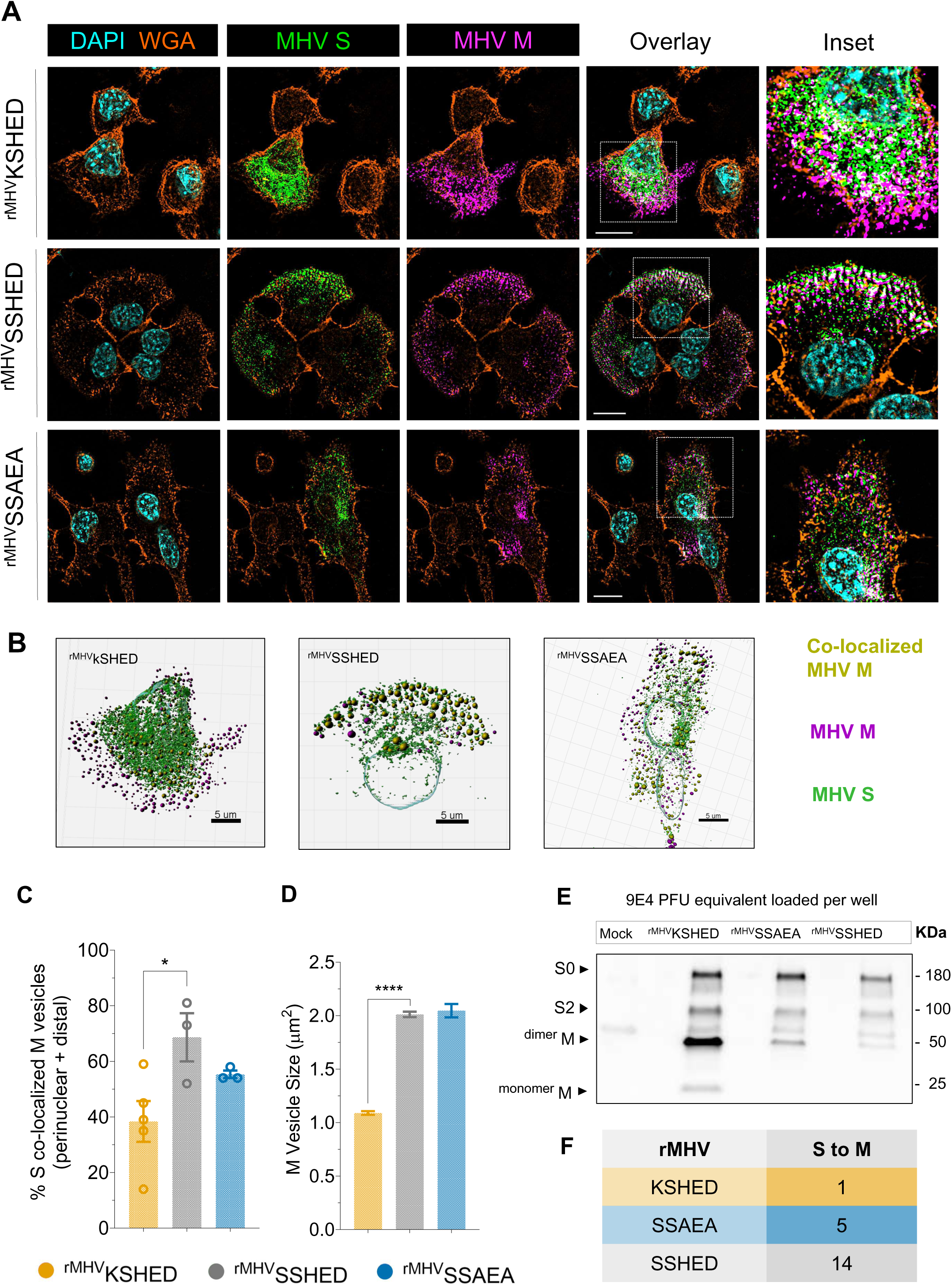
Coatomer-binding motifs dictate S–M interactions and virion composition. **(A)** KSHED, SSHED, SSAEA infected cells were co-stained for MHV S (in green) and M (in magenta) proteins. Data representative of three independent experiments. Scale bars, 10 µm. **(B)** 3D Iso-surface reconstruction of S surfaces and M spots for object-based co-localization analysis. M spots (magenta) within 200nm distance of a S surface (green) were deemed co-localized (yellow). Scale bars, 5 µm. **(C)** Quantification of the percentage of total M vesicles co-localized with S proteins, and **(D)** M vesicle size. Data graphed as mean ± SEM and analyzed using ordinary one-way ANOVA with Dunnett’s multiple comparisons test. **(E)** Comparative western blot analysis of PFU equivalent viruses for S and M proteins. **(F)** Densitometry analysis performed on protein bands to calculate S to M ratios.

Such localization changes were accompanied by altered M vesicle morphology. KSHED M vesicles were significantly smaller than in SSHED- or SSAEA-virus infected cells **(Fig. 5D)**. These observations may point to differential characteristics of KSHED egressing vesicles (marked by M), compared to SSHED or SSAEA. In addition to retrograde trafficking S molecules, COPI vesicle fusion to ERGIC is likely to expand the budding compartment. This may explain the smaller M vesicles size in the KSHED condition, where COPI vesicles are unable to deliver S molecules to ERGIC.

We next quantified S and M content in purified virions by western blot. Immunoblots revealed similar S levels across all three viruses, consistent with loading equivalent PFUs. Increased M protein signals were observed in KSHED and SSAEA virions compared to SSHED **(Fig. 5E).** Densitometric analysis of S-to-M signal ratios indicated that compared to SSHED virions, S densities on SSAEA and KSHED virions were ∼3- and 14fold lower, respectively **(Fig. 5F).**

Together, these data indicate that coatomer-binding motif alterations re-localize S proteins, reduce S–M interactions, and diminish S incorporation into virions, revealing coatomer engagement as a key determinant of virion composition and assembly.

#### Altered coatomer-binding affinity reorganizes S distribution and MHV particle architecture

The impact of modifying coatomer interactions on coronavirus morphology is not known. To assess whether mutations in the S coatomer-binding motif affect virion structure, we examined particle morphology and S organization by negative stain transmission electron microscopy (TEM) and cryo-electron tomography (cryo-ET).

Negative-stain TEM of glutaraldehyde-fixed virions revealed pronounced differences in peripheral S content. MHV-SSHED particles frequently carried ≥10 radially projecting S molecules, often in clusters, with only ∼10% of particles displaying fewer than five S molecules **(Fig. 6A, B)**. In contrast, ∼75% of MHV-SSAEA particles were sparsely decorated, and only ∼4% carried ≥10 S molecules, consistent with biochemical evidence of reduced incorporation **(Fig. 5E, 6B, C; SI Table T6)**. These results indicate that coatomer-binding affinity regulates both S incorporation and clustering on the virion surface.

**Figure 6.**
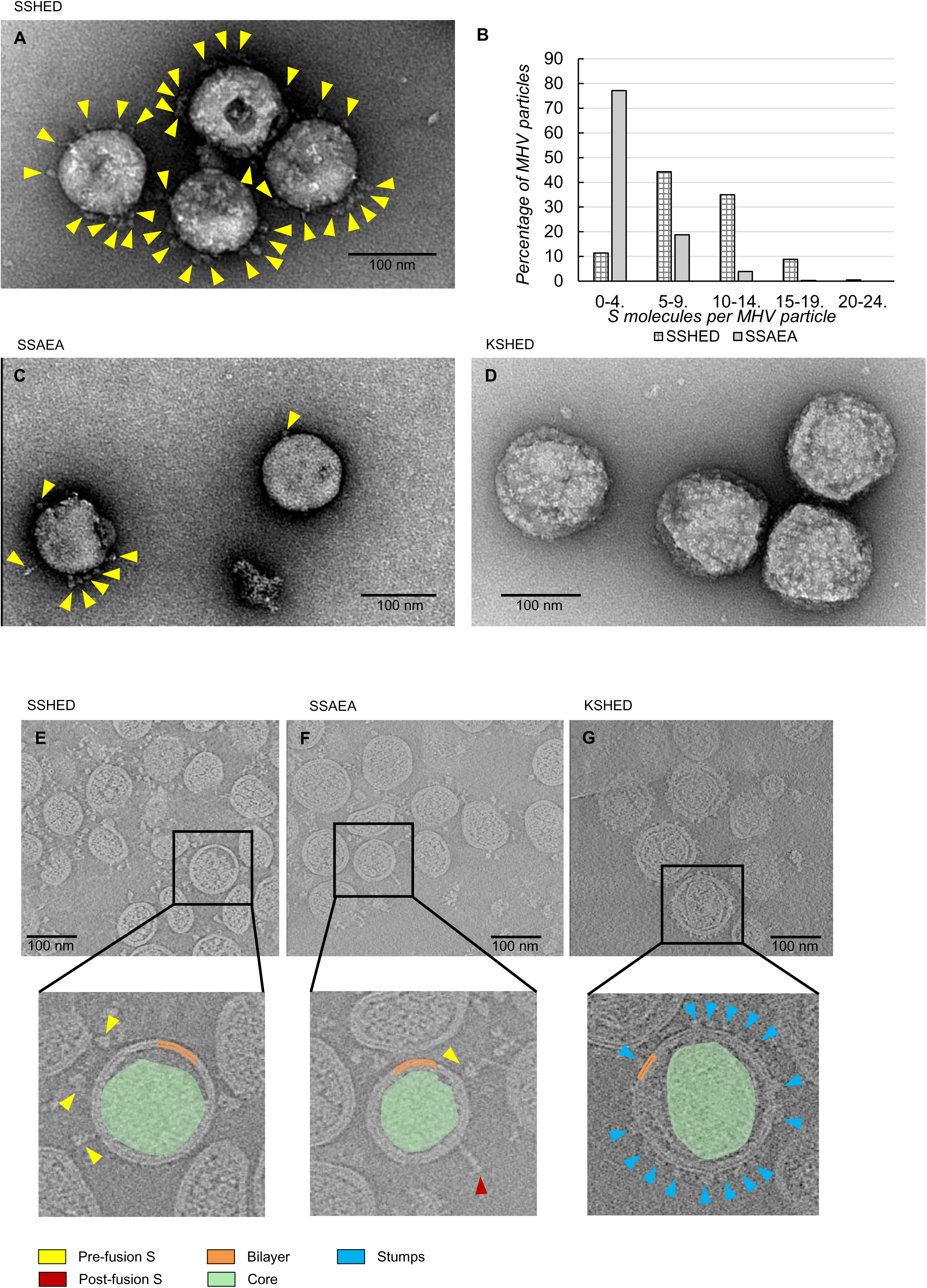
Altered coatomer-binding affinity reorganizes S distribution and MHV particle architecture. **(A)** Peripheral S clusters (yellow) in negatively stained MHV-SSHED particles. **(B)** Quantification of peripheral S clusters in negatively stained TEM images, showing increased abundance in MHV-SSHED compared to MHV-SSAEA. **(D)** MHV-KSHED particles exhibit a bumpy surface devoid of S molecules. **(E–F)** Cryo-ET reconstructions of MHV-SSHED **(E)** and MHV-SSAEA **(F)** reveal a bilayer with closely apposed core and surface S molecules. **(G)** MHV-KSHED particles display surface stumps and a detached core.

MHV-KSHED particles were morphologically heterogeneous exhibiting a distinct “bumpy” morphology, characterized by a uniform array of rectangular projections or “stumps” that lacked canonical S features **(Fig. 6D)**. The abundance of secreted M protein in KSHED-infected cultures supports a viral origin of these bumpy particles. Similar particles were occasionally observed in SSHED and SSAEA samples but at much lower frequency.

Cryo-ET of vitrified virions confirmed and extended these findings. SSHED and SSAEA particles displayed surface S molecules in prefusion conformations, with occasional post-fusion forms, overlying an internal bilayer consistent with the viral envelope and an electron-dense core attributed to RNA–nucleoprotein complexes **(Fig. 6E, F; SI Movies M5, M6)**. SSHED particles exhibited denser S clustering than SSAEA, matching TEM observations.

In contrast, bumpy KSHED particles showed a reorganized architecture consisting of two discrete zones: (i) an outer layer of lattice-like surface stumps contiguous with a bilayer, and (ii) a condensed, frequently polygonal core that was physically separated from the bilayer **(Fig. 6G; SI Movie M7)**. This structural uncoupling was not observed in SSHED or SSAEA particles, where the core remained closely associated with the bilayer.

Together, these analyses identify S–coatomer interactions as key determinants of coronavirus architecture. Mutations that strengthen coatomer binding drive virion heterogeneity and decouple the envelope from the nucleocapsid, likely contributing to reduced infectivity.

## DISCUSSION

Our investigation identifies an unconventional basic–acidic motif (HxD) in the coronavirus S tail as a critical regulator of MHV assembly, morphology, and infectivity. This motif directly occupies the WD40 binding site of the cellular coatomer, normally engaged by canonical clients, with intra-motif interactions priming it for coatomer recognition. Perturbing S–coatomer engagement disrupts secretory trafficking of both S and M proteins and drives the emergence of adaptive mutations in culture. These changes manifest as striking alterations in virion morphology, S organization, and infectivity, underscoring the central role of this motif in coronavirus biogenesis.

Our study reveals structural insight into the divergent evolution of the coatomer-binding motif. The canonical extended motif, defined by a basic–basic–acidic sequence, engages the αWD40 and β2WD40 domains through electrostatic complementarity^29,35–37^. Whereas prior work showed that coronavirus S proteins in SARS-CoV, SARS-CoV-2, and MERS-CoV mimic only the N-terminal dibasic portion of the extended motif^25,29,37,45,46^, we demonstrate that mimicry of the basic–acidic pair, as seen in MHV, is sufficient for coatomer binding. Strikingly, full mimicry of all three residues disrupts progeny assembly, morphology, and infectivity, underscoring the evolutionary pressure to limit mimicry and allow proper disengagement from coatomer. This framework helps explain how S proteins lacking the dibasic motif traffic in coronavirus infections.

Our study revises the prevailing model of coronavirus assembly, which has largely been shaped by VLP systems and co-expression studies^17,25,29^. Prior work suggested that S incorporation depends on M–S interactions within the Golgi network, with the coatomer delivering this complex to the assembly site^25,29^. However, our results challenge this Golgi-centric framework. If such a pathway operated during infection, loss of coatomer binding—as in MHV-SSAEA—would be expected to yield S-deficient, non-infectious virions. Instead, we find that MHV-SSAEA remains infectious although with reduced S content and markedly impaired S clustering. These observations suggest that M–S interactions are initiated earlier in the secretory pathway, likely within the ER or ERGIC, rather than in post-ERGIC Golgi compartments although this interaction likely continues in later compartments. This reinterpretation positions the early secretory system as the primary locus of coatomer-dependent assembly and reveals how coronaviruses leverage trafficking flexibility to maintain infectivity under host-imposed constraints. By shifting the conceptual framework from a static Golgi model to a dynamic, ER/ERGIC-centered process, our investigation sets the stage for future experiments to test whether M directly facilitates S export from the ER to the ERGIC, thus clarifying the spatial hierarchy of assembly events. Together, these findings refine the mechanistic map of coatomer-dependent coronavirus biogenesis and underscore the adaptability of viral membrane trafficking routes during infection.

The observation that coatomer interactions are not strictly required for infectious virion formation raises a key question: why do coronaviruses hijack the coatomer if S molecules can transit directly from the ER to assembly sites? Our structural studies reveal that coatomer primarily facilitates clustering, rather than delivery, of S molecules. Pre-assembled clusters of S, rather than individual trimers, appear preferentially trafficked during virion biogenesis, consistent with a coatomer-driven en bloc transfer mechanism. Such clustering would enhance infectivity, as only a subset of S tails needs to engage the coatomer to mobilize entire clusters. S palmitoylation in early secretory compartments^47–52^—the sites of coatomer activity—likely promotes this congregation. Clustering likely increases avidity for coatomer engagement, which is counter-balanced by limiting mimicry of the coatomer binding motif thus allowing controlled release at assembly sites. Coupled with the rate-limiting nature of trimerization^21^, this mechanism may act as a quality-control checkpoint, favoring incorporation of properly folded S trimers from the Golgi over misfolded or monomeric species from ER. Overall, these findings establish clustering as a central principle modulating coatomer interactions, with broader implications for oligomerization-dependent trafficking beyond viral biology.

Whereas the MHV-SSAEA virus revealed new roles for the coatomer in S clustering and clarified essential S–M interactions, the MHV-KSHED system highlights the opposite extreme—where excessive coatomer affinity drives viral adaptation. Unlike the infectious and genetically stable MHV-SSAEA, MHV-KSHED rapidly accumulates adaptive mutations, revealing how coronaviruses evolve under restrictive trafficking conditions. In MHV-KSHED populations, we identified S variants that either deleted the coatomer-binding motif or reverted to a canonical dibasic sequence, including a KxHxT tail also present in SARS-CoV-2. The comparable infectivity of the KxHxT mutant and wild type suggests that mimicry of either the dibasic or basic–acidic halves of the motif can equivalently support S delivery for assembly. Although VLP studies predicted that high-affinity KSHED S molecules would be excluded from assembly, analyses of infectious virions revealed a more flexible network: KSHED-bearing S proteins are incorporated—albeit at reduced levels—and promote secondary mutations in M, E, and possibly NS2a that facilitate S extraction from coatomer. These findings identify S as a central hub for both assembly and adaptability, positioning coatomer–S interactions as key evolutionary constraints in coronavirus biogenesis.

Our investigation also provides structural insight into the consequences of enhanced coatomer–S interactions, a feature previously unexplored. Based on the low S content in MHV-KSHED particles, one might have expected a morphology similar to MHV-SSAEA, characterized by smooth, sparsely decorated surfaces. However, electron microscopy reveals a strikingly different architecture: MHV-KSHED virions display polygonal cores and a lattice-like, bumpy surface devoid of normal S decoration. The emergence of these previously unreported bumpy particles identifies coatomer–S affinity as a key determinant of coronavirus morphology. This finding broadens the structural framework of coronavirus biogenesis and suggests that even single-residue alterations in coatomer–S interactions can reprogram virion assembly. Future characterization of their protein and nucleic acid composition will be critical to determine whether these bumpy particles represent malformed intermediates, alternative assembly states, or adaptive structural variants.

Our findings highlight how coronaviruses limit coatomer–S interactions to preserve host secretory trafficking. High-affinity KSHED S molecules redistributed COPB2 from distal compartments to perinuclear regions, suggesting that strong S engagement traps COPI vesicles in early secretory cycling. Such relocalization has broad implications: it may restrict retrograde transport of endogenous cargo, alter the composition of the secretory proteome, and disturb membrane homeostasis. In the extreme, these disruptions could compromise cellular survival and function. By contrast, wild-type SSHED S molecules preserved relatively normal COPB2 distribution, implying that coronaviruses have evolved a finely tuned level of coatomer engagement that permits efficient assembly while preserving the host trafficking processes needed to form and maintain virus replication organelles. This balance underscores a central principle of host–virus adaptation: viral proteins must co-opt essential host machinery without destabilizing its core functions. Defining how coronavirus infection rewires coatomer biology at the systems level will be essential to understand the broader cellular consequences of S–coatomer engagement.

Beyond virology, these results expand the coatomer recognition code itself. By establishing that basic–acidic residues can substitute for a canonical dibasic motif^33,35,36^, our work points to new classes of endogenous coatomer clients and provides a foundation for discovering alternative modes of coatomer trafficking. More broadly, our investigation defines a fundamental organizing principle of coatomer-binding motifs: functional engagement requires occupation of the –3 position and one additional site—either –5 or –1—by residues compatible with the interaction. The –3 position emerges as indispensable, as it dictates the main-chain conformation necessary for productive binding. Testing such hypotheses in BLI binding assays was enabled by stabilizing the often intractable αWD40 domain^35,36,42^ using Fab17, which opens the door to such high-resolution structural and functional studies of coatomer–client interactions, providing a broadly applicable tool for dissecting coatomer biology.

In conclusion, our findings reveal that precisely tuned coatomer–S engagement, mediated by the unconventional HxD motif, controls coronavirus assembly, virion morphology, and infectivity, while also uncovering a new class of uncharacterized ‘bumpy’ particles for future structural and functional exploration.

## Supporting information

SI Move M5

## Abbreviations

αWD40: N-terminal WD40 domain of coatomer α subunit
β2WD40 or β΄WD40: N-terminal WD40 domain of coatomer β2 or β΄ subunit
COPB2: coatomer β2 subunit
CPER: circularized polymerase extension reaction
cryo-ET: cryogenic electron tomography
E: envelope protein
ER: endoplasmic reticulum
ERGIC: endoplasmic reticulum–Golgi intermediate compartment
GEQ: genome equivalents
M: membrane protein
MHV: murine hepatitis virus
N: nucleocapsid protein
PFU: plaque-forming unit
RNP: ribonucleoprotein
S: spike
TEM: transmission electron microscopy

## SUPPLEMENTARY INFORMATION

### Figures

S1 | HxD motif in the MHV S tail mediates selective binding to αWD40-Protein production and characterization.

S2 | HxD motif engages distinct binding environments in αWD40 and β2WD40 domains-MD simulations.

S3 | S-coatomer affinity governs MHV infectivity and genetic adaptation-Engineering recombinant MHV-A59 viruses with altered S-coatomer affinity

S4 | Coatomer association controls plasma membrane S distribution and syncytia formation

### Tables

T1 | Sequence relationships between coatomer WD40 domains

T2 | Peptides used in this investigation

T3 | BLI assays of MHV S tail peptides with coatomer WD40 domains

T4 | Crystallographic data collection and refinement statistics

T5 | Genetic adaptation by rMHVs with altered S-coatomer affinity

T6 | Coatomer-dependent modification of S content in MHV particles imaged by negative staining TEM

### Movies

M1 | Morph animation shows conformational change in S tail His1322 side-chain from αWD40 to β2WD40 domain bound states. The red sphere represent a crystallographically ordered water molecule in the β2WD40 co-crystal structure.

M2 | Coatomer association controls subcellular distribution of KSHED S molecules

M3 | Coatomer association controls subcellular distribution of SSHED S molecules

M4 | Coatomer association controls subcellular distribution of SSAEA S molecules

M5 | Altered coatomer-binding affinity reorganizes S distribution and MHV particle architecture- Cryo-ET reconstruction of purified MHV-SSHED particles

M6 | Altered coatomer-binding affinity reorganizes S distribution and MHV particle architecture- Cryo-ET reconstruction of purified MHV-SSAEA particles

M7 | Altered coatomer-binding affinity reorganizes S distribution and MHV particle architecture-Cryo-ET reconstruction of purified MHV-KSHED particles

## ACKNOWLEDGEMENTS

Research reported in this publication was supported by National Institutes of Health grants from National Institute of General Medical Sciences (NIGMS) R01GM150187 (SSH) and National Institute of Allergy and Infectious Diseases (NIAID) R21AI176252 (TG), and an Arthur J. Schmitt Leadership Scholars Dissertation Fellowship (SM). The content is solely the responsibility of the authors and does not necessarily represent the official views of the National Institutes of Health. We would like to thank Drs. Ed Campbell, Adarsh Dharan, and David J. Rademacher (Loyola University) for their support with Lattice SIM; Dr. Edwin Pozharski (University of Maryland School of Medicine) for assistance with electron microscopy; Dr. Duncan Souza, Dr. Kai Cai, and Ms. Barbara Smith (Johns Hopkins University) for assistance with electron microscopy; Dr. Jean Jakoncic (Brookhaven National Lab) for X-ray diffraction data collection and processing; Mr. Nevin Brittain (University of Chicago) for Fab production. A Shared Instrumentation Grant from the National Institutes of Health (1S10OD034431-01) supported this work. This article was supported by funds through the Maryland Department of Health’s Cigarette Restitution Fund Program – CH-649-CRF and the National Cancer Institute - Cancer Center Support Grant (CCSG) – P30CA134274. This research used a Thermo Fisher Scientific Mark IV and computational resources at the Maryland Center for Advanced Molecular Analysis (M-CAMA). This research used resources at the 17-ID-1 AMX (Highly Automated Macromolecular Crystallography) of the National Synchrotron Light Source II, a U.S. Department of Energy (DOE) Office of Science User Facility operated for the DOE Office of Science by Brookhaven National Laboratory under Contract No. DE-SC0012704. The Center for BioMolecular Structure (CBMS) is primarily supported by the National Institutes of Health, National Institute of General Medical Sciences (NIGMS) through a Center Core P30 Grant (P30GM133893), and by the DOE Office of Biological and Environmental Research (KP1605010).

## AUTHOR CONTRIBUTIONS

Conceptualization: S.S.H., T.G.; Methodology: S.M., A.K.S., S.S., F.L.K., T.K., S.E., D.D., A.K., T.G., S.S.H.; Software: F.L.K.; Validation: S.M., A.K.S., S.S., F.L.K., T.K., S.E.; Formal analysis: S.M., S.S., A.K.S, F.L.K., K.J., D.D., T.G., S.S.H.; Investigation: S.M., S.S., A.K.S., F.L.K., D.D., B.J., B.D., T.K., T.G., S.S.H.; Resources: T.K., A.K., R.E.A.,T.G., S.S.H.; Data curation: S.M., S.S., A.K.S., T.K., T.G., S.S.H.; Writing original draft: S.M., A.K.S., S.S., F.L.K., T.K., S.E., A.K., R.E.A., T.G., S.S.H.; Writing - Review & Editing: All authors; Visualization: S.M., A.K.S., S.S., F.L.K., T.B., S.S.H.; Supervision: R.E.A., A.K., T.G., S.S.H.; Project administration: T.G., S.S.H.; Funding acquisition: T.G., S.S.H.

## DISCLAIMER STATEMENT

During the preparation of this work the authors used ChatGPT and Grammarly to improve readability. After using these tools, the authors reviewed and edited the content as needed and take full responsibility for the content of the published article.

