## Supplementary material for "An unconventional HxD motif orchestrates coatomer-dependent coronavirus morphogenesis": Materials and Methods

**Cell Lines**

BL21(DE3)pLysS cells from Promega were cultivated in 2XYT broth (Fisher Scientific). Expi293 cells from Thermo Fisher Scientific (TFS) were maintained in Expi293CD media at 70% humidity and 8% CO2. Human embryonic kidney 293T cells, kindly provided by Dr. Edward Campbell (Loyola University Chicago), were maintained in Dulbecco’s Modified Eagle Media (DMEM, Corning) supplemented with 10% heat-inactivated Fetal Bovine Serum (FBS, Bio-Techne), 100 U/ml Penicillin G, 100 µg/ml streptomycin, 0.1 mM non-essential amino acid, 100 nM sodium pyruvate, 10 mM HEPES (pH 7.3). Murine delayed brain tumor cells (mDBT)1 and murine 17CL1L2 fibroblasts were generously provided by Dr. Susan Baker (Loyola University Chicago). mDBT cells were cultured in Minimal Essential Media (MEM, Corning) supplemented with 5% FBS, 10% tryptose phosphate broth (homemade), 100 U/ml penicillin G, 100 µg/ml streptomycin, 26.8 mM sodium bicarbonate, and 2 mM L-glutamine. 17CL1 cells were cultured in DMEM media supplemented with 5% FBS, 5% tryptose phosphate broth, with 2 mM L-Glutamine, 100 U/ml penicillin G, 100 µg/ml streptomycin. HEK293T, mDBT, and 17CL1 cell lines were maintained at 37°C and 5% CO2.

**Protein expression and purification**

Recombinant production of S. pombe αWD40 (wild type) and mutant (His31Ala) were carried out in E. coli as published previously^1^ with two modifications- first, the affinity elution buffer was supplemented with 0.02% (weight/volume) of the detergent, n-docedyl-β-D-maltopyranoside (DDM, Anatrace), and protease cleavage was performed with an equimolar ratio of in-house purified ULP1 enzyme at 4 °C for 1-1.5 hours. Mouse β2WD40 domain was sub-cloned into a pcDNA3.1(+) vector with an N-terminal extension encoding a twin-strep tag, Hisx6 tag, and SUMO-star tag. Transection in Expi293 cells was performed using performed using polyethyleneimine 25k, followed by cell culture and purification as described. This construct is stable and does not require detergent or glycerol during cleavage with SUMO-star protease (Enzo Life Science). After removal of the cleaved tag on a Ni-NTA column, size exclusion chromatography (SEC) was performed on either a Superdex 200 increase 10/300 GL column or Superdex 200 pg 16/600 chromatography column (Cytiva) pre-equilibrated with buffer 20 mM Tris-HCL pH 7.5, 150 mM NaCl and 200 µM TCEP. Fab17 production was performed as published elsewhere^2^.

**Biolayer interferometry**

Biotinylated MHV S-tail peptides (Biomatik USA) were immobilized on streptavidin biosensors (Sartorius) at 4 µg/ml in kinetics buffer (100 mM HEPES (pH 7.5), 150 mM NaCl, 0.5% BSA, and 0.02% Tween). Binding kinetics were determined using the αWD40-Fab17 complex at concentrations of 5, 2.5, 1.25, and 0.675 µM. For non-binder peptides, assays were performed using a single αWD40-Fab17 and β2WD40 concentration at 5 µM. All measurements were performed in triplicate using an Octet RED96e system (Sartorius). Data were reference-subtracted and globally fitted to a 1:1 binding model using the Octet Data Analysis software (version 11.1, Sartorius).

**X-ray crystallography**

Yeast αWD40 domain concentrated to 2.5 mg/ml and incubated in incubated with the MHV S tail heptapeptide (NISSHED, in DMSO) in the molar ratio of 1:2 for 20 minutes. Crystallization screening was performed at 295 K on a Mosquito LV crystal robotic system (SPT LabTech) using 96-well sitting drop plates by mixing 0.1 µL of αWD40-peptide complex with 0.1 µL of precipitant and various commercially available screens (Crystal screen HT, PEGRx HT, PEG/Ion HT, Index HT, JCSG Plus HT and Wizard classic from Hampton Research and Rigaku). One of the conditions from JCSG Plus HT (Condition B4) yielded crystals in 2 days. Rod shape co-crystal of αWD40 domain was obtained in 0.1 M HEPES pH 7.5, 10% w/v PEG 8000 and 8% v/v ethylene glycol. Crystallization experiment of αWD40 domain mutant (H31A) was set using 24 well hanging drop plate by mixing 1 µL protein with 1 µL reservoir. Rod shape crystals were obtained in 0.2 M sodium citrate tribasic dihydrate, 15% PEG 3350 at room temperature. Co-crystallization of β2WD40 domain (21 mg/ml) with the heptapeptide was performed as described previously, and crystals were obtained in multiple conditions such as 19-23% PEG 4000 with MES buffer (0.1 M, pH 6.2); 15-21% PEG 20,000 with MES buffer (0.1 M, pH 6.5); and 20-24% PEG 3350 with NaSCN (0.2 M). Single crystals were mounted on cryo-loops using 20% (volume/volume) glycerol as a cryo-protectant and flash-frozen in liquid nitrogen.

X-ray diffraction data were collected at the Brookhaven National Laboratory, National Synchrotron Light Source II (NSLS II) beamline 17-ID-1 AMX^3^. The data were processed using XDS in Dimple, the automated data acquisition and processing pipeline at the beamline^3,4^. Molecular replacement was performed using an apo-structure of the αWD40 domain (PDB ID 7S22)^5^ for αWD40-peptide complex and (PDB ID 8ENS)^1^ for the αWD40 H31A mutant. A homology model of the murine β2WD40 domain from SWISS-MODEL^6^ was used for molecular replacement in the β2WD40-peptide complex. Refinement and model building were performed in Phenix.refine^7,8^ and Coot^9^.

**MD simulations**

Rosettas BuildPeptide tool^10^ was used to build peptide structures of the four 21-mer sequences, VMD^11^ was then used to solvate each peptide in a TIP3P water box of size 100 Å x 100 Å x 100 Å and neutralized with 150 mM NaCl. All simulations were performed in three replicas with NAMD3^12,13^ and parameters described by CHARMM36m^14,15^ all-atom force fields using San Diego Supercomputing (SDSC) resources. Periodic Boundary Conditions^16^ were applied, Particle Mesh Ewald^17^ was used to calculate long-range Coulombic potentials, Lennard-Jones potentials^18^ were calculated according to a 10 Å – 12 Å cutoff and switching scheme, pairwise list generation was cutoff at 15.5 Å. Langevin thermostat and piston were used to maintain temperature at 310 K unless otherwise stated and at 1 atmosphere of pressure.

Minimization: To relax all conformations and alleviate close contacts resultant from CHARMM-GUI O-glycosylation steps, all solvated and neutralized structures were subjected to 10,000 steps of conjugate gradient energy minimization with no atomic restraints or constraints.

Heating: Systems were then heated from 10 K to 310 K, increasing temperature by 25 K every 10,080 steps, timestep 2 fs, for a total of 240 ps of heating. Once at 310 K, an additional short equilibration of 770 ps of simulation was conducted.

Equilibration: Mini1 and Mini2 were then subjected to NpT equilibration at 310 K for 0.5 ns, 2 fs timestep (useFlexibleCell option “on”, wrapWater option “on”, wrapAll option “on”). Systems were then subjected to an additional 55 ns NVT equilibration with 2 fs timestep with fixed box dimensions (useFlexibleCell option “off”).

Production: 550 ns of NVT initial conventional MD simulation production runs were conducted at 310 K, timestep 2 fs, saving frames every 5 ps. Following the above construction and simulation protocols, we then conducted extensive analyses of these simulations using MDAnalysis and in-house scripts^19,20^.

**Negative staining transmission electron microscopy**

Glutaraldehyde-fixed MHV virions were buffer exchanges into phosphate-buffered saline (PBS) using centrifugal filter units. Samples (5 µL) were applied onto formvar/carbon-coated copper grids (400 mesh) that had been glow-discharged for 30 s. After adsorption of the virus sample for 1 minute, the grids were edge-blotted, washed briefly with PBS, and negatively stained with 1% (w/v) uranyl acetate for 30 s. Excess stain was removed by edge-blotting and a vacuum, and the grids were air-dried. Imaging was performed at Johns Hopkins University Microscopy Facility on a TFS Talos L120C G2 Transmission Electron Microscope equipped with a TFS Ceta camera (cooled 16 Mpixel CMOS, 16-bit 1-25fps) at a magnification of 73,000 X corresponding to a pixel size of 1.908 Å. CTF estimation and manual particle picking were performed in EMAN2^21^. The analysis of peripheral S protein molecules was performed manually and individually for each particle.

**Cryo-electron tomography**

Glutaraldehyde-treated MHV virions were buffer exchanged into a final volume of approximately 50 µL in PBS using Amicon concentrators (100 kDa molecular weight cut-off; 0.5 ml volume). These samples were mixed in 1:4 beads:virus sample volumetric ratio with the fiducial marker, protein A gold (10 nm, Aurion). 3–4 µL aliquots were applied onto copper grids with ultrathin carbon support films (200 mesh, 2/2 spacing; Quantifoil). Grids were glow-discharged at 15 mA for 10 seconds prior to vitrification. Excess liquid was blotted for 4-5 s, and grids were plunge-frozen into liquid ethane cooled by liquid nitrogen using a TFS Vitrobot Mark IV under controlled temperature and humidity conditions. Grids were stored in liquid nitrogen until imaging. Screening for virus and fiducial marker distribution and proper ice thickness was performed at Johns Hopkins University on a TFS Glacios Microscope equipped with a Falcon 4i direct electron detector. The selected grids were then shipped under cryogenic conditions to Purdue University for data collection on a TFS 300 kV Titan Krios G4 TEM equipped with a Gatan K3 DED and a Gatan Quantum GIF energy filter. Tilts were acquired at a nominal magnification of 64,000X resulting in a super-resolution pixel size of 0.666Å/pixel and an exposure time of 259ms split over 10 frames, resulting in a total dose of 2.91e/Å2 per tilt. All tilt series were recorded in a dose symmetric fashion from -60˚ to +60˚ in 3˚ increments and a grouping of 2. Data processing for motion correction, CTF estimation, and tilt series alignment were performed in Warp^22^ and tomograms were binned to 10Å/pixel for visualization.

**Virus engineering and sequencing**

MHV strain A59 was propagated in mDBT cells. Recombinant MHV-A59 viruses with altered coatomer binding motifs were generated using the circular polymerase extension reaction (CPER)^23,24^. Briefly, the MHV A59 genome was divided into 10 overlapping fragments. Fragment 8 contained the S gene. Following viral RNA extraction from infected cells, cDNAs fragments were synthesized using the SuperScript IV One-Step RT-PCR System (Invitrogen). Fragment 8 cDNA with intended S mutation(s) was generated via site directed mutagenesis and Single Tube Overlap Extension (SOE-ing) PCR. All 10 fragments and a UTR-linker (3’ overlap-HDV ribozyme-SV40 poly A signal-CMV promoter-5’ overlap) were mixed in equimolar amounts (0.1 pM/fragment) in a total reaction volume of 50 µL, with dNTPs, Prime Star GXL DNA polymerase, and 1X PS GXL reaction buffer. Initial denaturation was performed at 98 °C for 2 minutes, followed by 30 cycles of denaturation at 98 °C for 10 seconds, annealing at 60 °C for 15 s, extension at 68 °C for 15 min, and a final extension at 68 °C for 15 min. Circularized CPER products were stored at room temperature overnight before being transfected into HEK293T cells. At 1 day post-transfection, the transfected cells were co-cultured with mDBT cells that expressed the MHV receptor, murine CEACAM1. The co-culture was replenished with fresh mDBT and HEK293T media and monitored for virus induced cytopathic effects (CPE) for successful virus recovery.

For virus sequencing, intracellular RNA was harvested from producer and/or infected cultures. Extracted RNA served as template for cDNA synthesis across the structural genes. The resulting cDNA samples were sequence verified by Plasmidsaurus.

**Production of VLPs**

VLPs were produced in HEK293T cells as described previously^25,26^. Briefly, plasmids expressing the MHV A59 structural genes were mixed in a 1:1 (weight/weight) ratio and combined with LipoD293™ In Vitro DNA Transfection Reagent (SignaGen) and incubated with the cells for 6 to 8 hours. At 6-8 hours post incubation, transfection reagents were removed and the monolayer washed once with 1X PBS. The wells were replenished with complete media. VLPs secreted into the media were harvested between 20-30 hours post transfection.

**Virus and VLP Concentration**

Virus or VLP containing supernatants were first clarified via differential centrifugation at 4°C; first at 300 x g for 10 minutes, followed by 10 more minutes at 3000 x g. Clarified supernatant samples were carefully decanted to exclude cell debris pelleted at the bottom of the tube. Next, a 25% (weight/weight) sucrose cushion in serum free complete DMEM was carefully layered under the clarified samples, and centrifuged at 5000 x g at 4°C for ~20 hours. The pellet was resuspended in 2X serum-free media. In some cases, the virus pellet was further purified by ultracentrifugation using a discontinuous density gradient- 35%, 30%, 25%, 20%, and 15% (w/v)- of iodixanol (OptiPrepTM, STEMCELL Technologies) The virus containing sample in 40% (weight/volume) iodixanol was carefully layered at the bottom of the gradient containing tube. Samples were then centrifuged using an SW60 rotor, at 55,000 rpm for 2 hours at 4°C. Following ultracentrifugation, viruses were carefully fractionated into 200 uL samples.

**Virus Inactivation**

The virus samples were mixed with 1% glutaraldehyde at a 1:1 ratio (volume/volume) and incubated at room temperature for 1 hour. Virus inactivation was further validated using plaque assays.

**Plaque Assay**

Supernatant samples containing infectious viruses were 10-fold serially diluted and incubated with ~80% confluent mDBT monolayer in a 6 well plate. Plates were rocked every 15 minutes to ensure even distribution of incubating virus sample. At 2 hours post incubation, the infectious sample was removed and the monolayer washed once with 1X PBS. A 0.8% agarose overlay in 2X mDBT media was carefully applied to the monolayer. Once the agarose had solidified, the plaque assay plates were incubated for 48 hours. At the end of the infection, indicator mDBT cells were fixed with 3.7% formaldehyde in 1X PBS for 1 hour at room temperature and stained with 0.1% crystal violet solution. Plaque size was measured using the ViralPlaque macro on ImageJ.

**Viral RNA extraction**

TRIzolTM (Invitrogen) RNA extraction was performed for viral cDNA synthesis. For total intracellular RNA extraction, appropriate amount of TRIzol reagent was added to the cell monolayer (0.3 mL/million cells) followed by homogenization. Molecular grade chloroform was added to the TRIzol homogenate at a volume/volume ratio of 1:3, and centrifuged at 12,000 x g at 4◦C for 10 minutes. RNA was carefully harvested from the aqueous layer, transferred to a clean microfuge tube, and precipitated using 5 µg (total) Glycogen and 100% Isopropanol (at a weight/volume ratio of 1:1) followed by centrifugation at 12,000 x g at 4◦C for 1 hour. The RNA pellet was washed in 75% ethanol, air-dried, and resuspended in appropriate volume of RNAse free dH2O.

Automated KingFisher™ nucleic acid extraction was performed to harvest intracellular and extracellular RNA for infection kinetics experiment. KingFisher instrumentation and reagent support was kindly provided by Dr. Susan Uprichard (Loyola University Chicago). KingFisher MagMax Pathogen Kit (cat# 4462359) was used to harvest extracellular RNA, and the CORE Kit (cat# A32702) was utilized for intracellular RNA extraction.

**Quantitative RT-PCR**

A primer set corresponding to a 75 base-pair region on CoV-nsp4 was used in qRT-PCR. Experiments were performed using the Luna Universal One-Step RT-qPCR Kit (New England Biolabs) on the CFX Opus 96 Real-Time PCR System (Bio-Rad). Data was reported as genome copies/equivalence/mL. Briefly, samples containing known copies of the target amplicon were serially diluted (10-fold) to generate a standard curve. The standard curve was included with every run and used to extrapolate genome copy numbers for the samples analyzed. For intracellular samples, cycle threshold (ct) values were also normalized to the housekeeping gene GAPDH.

**Western blotting**

Samples were mixed with 1X Laemmli buffer (2% weight/volume SDS, 0.06 M Tris-HCl, 10% glycerol, 0.01% bromophenol blue, and 2% β-mercaptoethanol (BME) and denatured by heating at 95˚C for 10 minutes. Denatured samples were electrophoresed through 4-20% Mini-PROTEAN TGX Precast Protein Gels (Bio-Rad). Following electrophoresis, proteins were transferred to nitrocellulose membranes (Bio-Rad) using the semi-dry transfer method on a Trans-Blot® TurboTM system (Bio-Rad). Nitrocellulose membranes with proteins of interest were blocked at room temperature for 1 hour using 5% milk in 1X TBST (25 mM Tris-HCl, 140 mM NaCl, 27 mM KCl, and 0.05% Tween 20), and incubated with primary antibodies (in 1% milk in 1X TBST) overnight at 4◦C. The membranes were subsequently incubated with appropriate secondary antibodies (conjugated to horseradish peroxidase) for 1 hour at room temperature. Finally, the membranes were incubated with a chemiluminescent substrate (Pierce™ ECL / SuperSignal™ West Femto from Thermo Scientific) and imaged on a FluorChem E System (92-14860-00) by ProteinSimple.

**Immunofluorescence staining**

For immunofluorescence microscopy, cells were seeded onto fibronectin coated #1.5 coverslips. At the end of infection, coverslips were fixed with 4% paraformaldehyde and washed with 1X PBS. Cells used for intracellular staining were permeabilized with 0.1% Triton X 100. Subsequently, the coverslips were blocked in blocking buffer (10% goat-serum in 1X PBS) overnight at 4◦C. Primary and secondary antibody incubations were performed for 1 hour and coverslips were mounted onto glass slides using the SlowFade™ glass soft-set antifade mountant (Invitrogen).

**Fluorescence microscopy**

Super-resolution structured illumination microcopy was performed on ZEISS Lattice SIM 5, equipped with Hamamatsu-ORCA Fusion BT cameras. Z stacks at 5µm depth were captured at 0.1 µm intervals using the lattice SIM 3D leap mode with a 63X oil-immersion objective. SIM² image reconstruction was performed following the recommended settings for fixed samples. Identical conditions were applied to all image acquisitions.

**Fluorescence microscopy image analysis**

All images were analyzed using Bitplane Imaris 9.9 (Oxford Instruments). For 3D iso-surface reconstruction, the spot and surface creation features were used. Additionally, distances between objects were calculated using the “shortest distance calculation” feature. For object-based co-localization analysis, 3D objects within a distance of 200 nm were considered co-localized.

**Statistical Analysis**

GraphPad Prism was used to perform all statistical analyses. Data were plotted with error bars including the standard error mean (SEM). Statistical tests used include one-way/two-way ANOVA, unpaired students t test. Statistical significance was concluded if p values were less than 0.05.
