## Supplementary material for "An unconventional HxD motif orchestrates coatomer-dependent coronavirus morphogenesis": SI Fig. S1

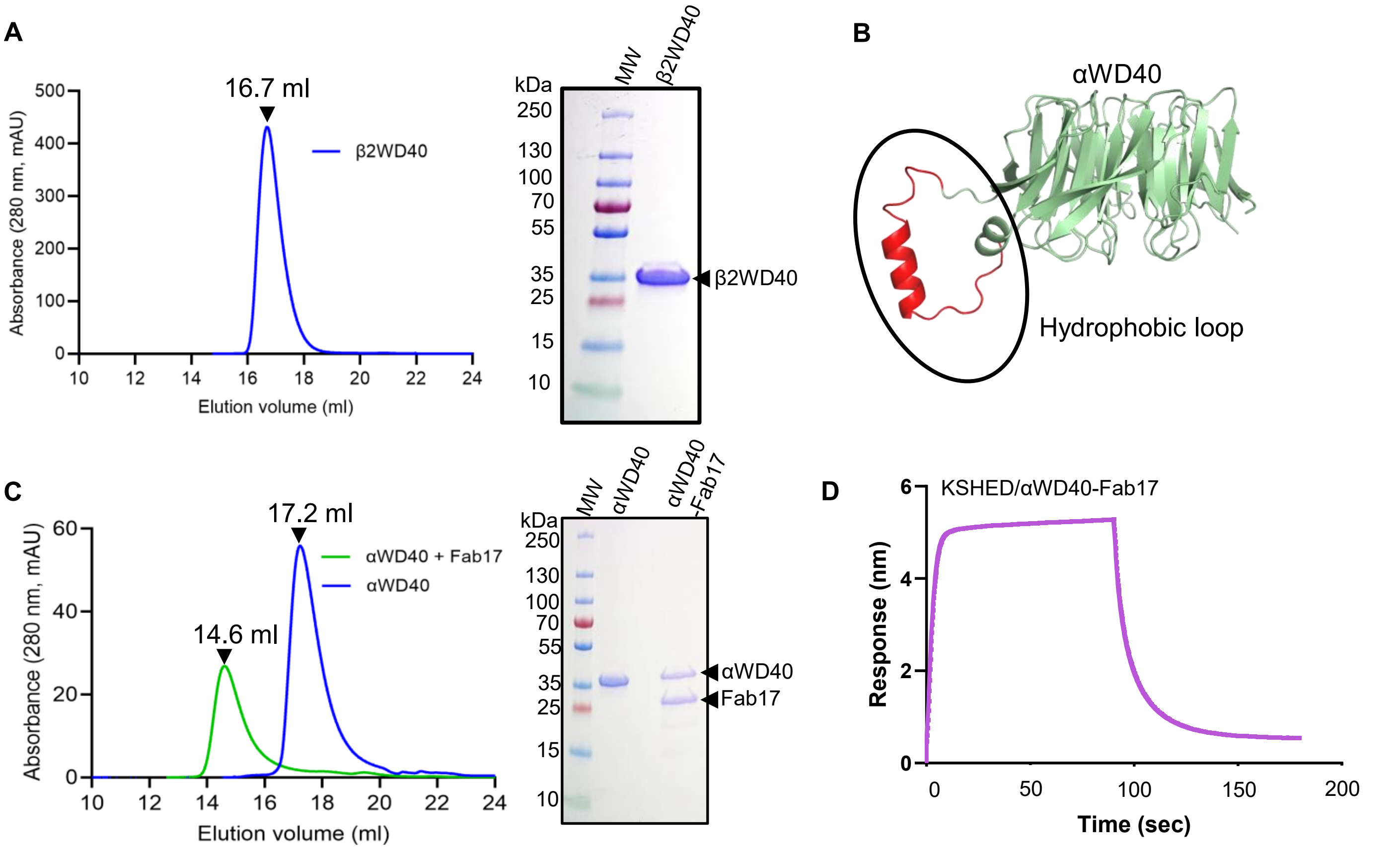


**SI Fig. S1 | HxD motif in the MHV S tail mediates selective binding to αWD40- Protein production and characterization. (A)** Size-exclusion chromatography (Superdex 200 Increase 10/300) and SDS-PAGE analysis show the purified murine β2WD40 domain. **(B)** Hydrophobic loop on the surface of αWD40 domain. **(C)** Size-exclusion chromatography (Superdex 200 Increase 10/300) and SDS-PAGE analysis show the purified murine αWD40 domain and the αWD40–Fab17 complex. **(D)** BLI sensorgram shows enhanced response (nm) for KSHED 21-mer peptide to αWD40–Fab17 (5 μM) relative to SSHED 21-mer in Figure 1.
