## Supplementary material for "An unconventional HxD motif orchestrates coatomer-dependent coronavirus morphogenesis": SI Fig. S2

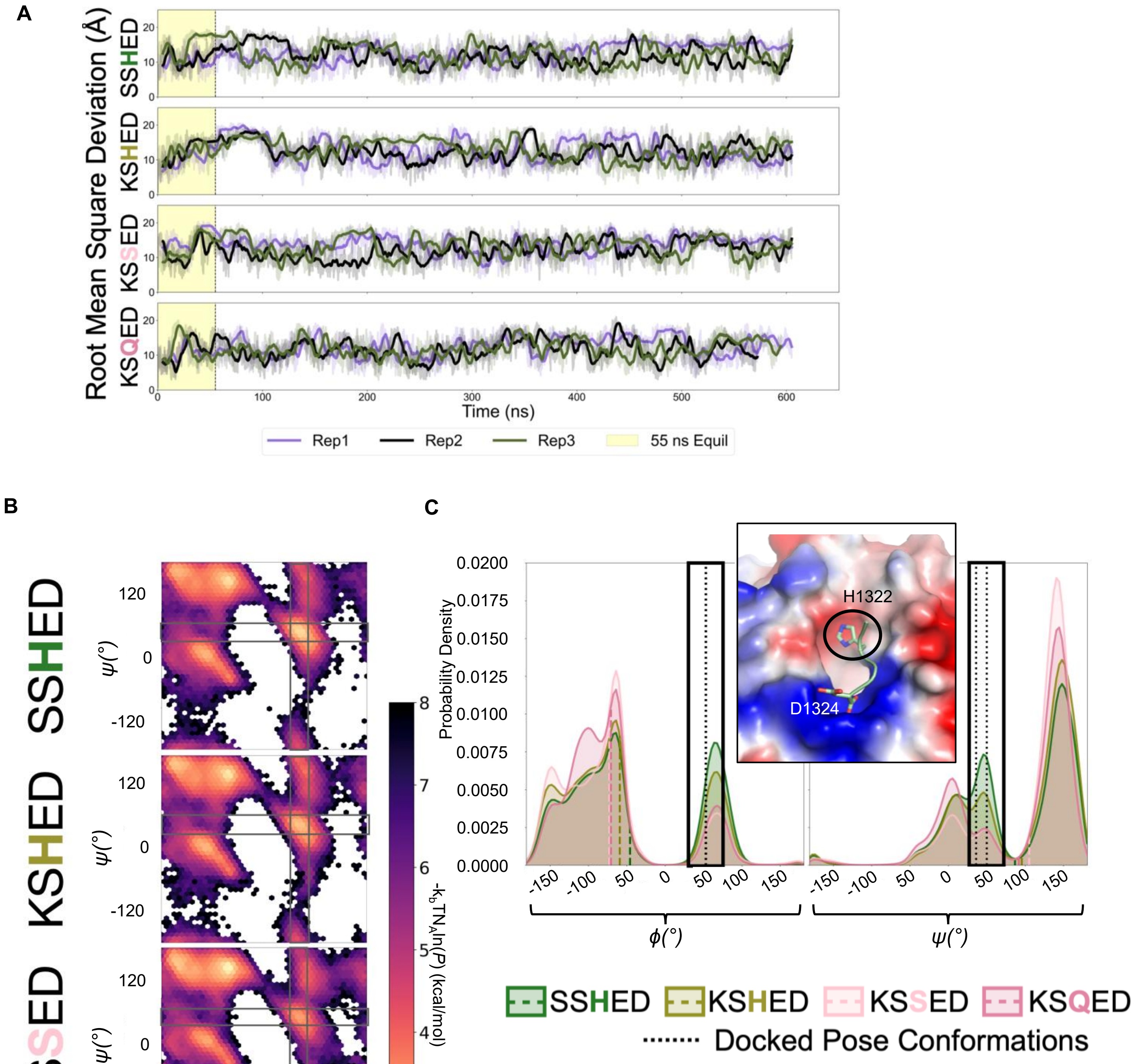


**SI Fig. S2 | HxD motif engages distinct binding environments in αWD40 and β2WD40 domains- MD simulations. (A)** Root Mean Square Deviations calculated for each peptide and each frame over the course of 55 ns of equilibration simulation conditions and then 550 ns of production simulations. All trends reveal well equilibrated and stable MD simulations, thus minimizing the possibility of drifting or unequilibrated data. **(B)** Ramachandran plots of the –3 residue from 550-ns production simulations of 21-mer peptides ending in SSHED, KSHED, KSSED, or KSQED. Grey boxes mark the Φ/Ψ ranges observed in docked poses. **(C)** One-dimensional Φ/Ψ kernel probability densities for the –3 residue in SSHED (green), KSHED (moss green), KSSED (light pink), and KSQED (pink). Dashed lines denote distribution means; dotted black lines indicate dihedrals from two docked poses. The black box defines the torsional range considered compatible with binding. The inset highlights His1322 in the MHV tail peptide.
