## Supplementary material for "An unconventional HxD motif orchestrates coatomer-dependent coronavirus morphogenesis": SI Fig. S3

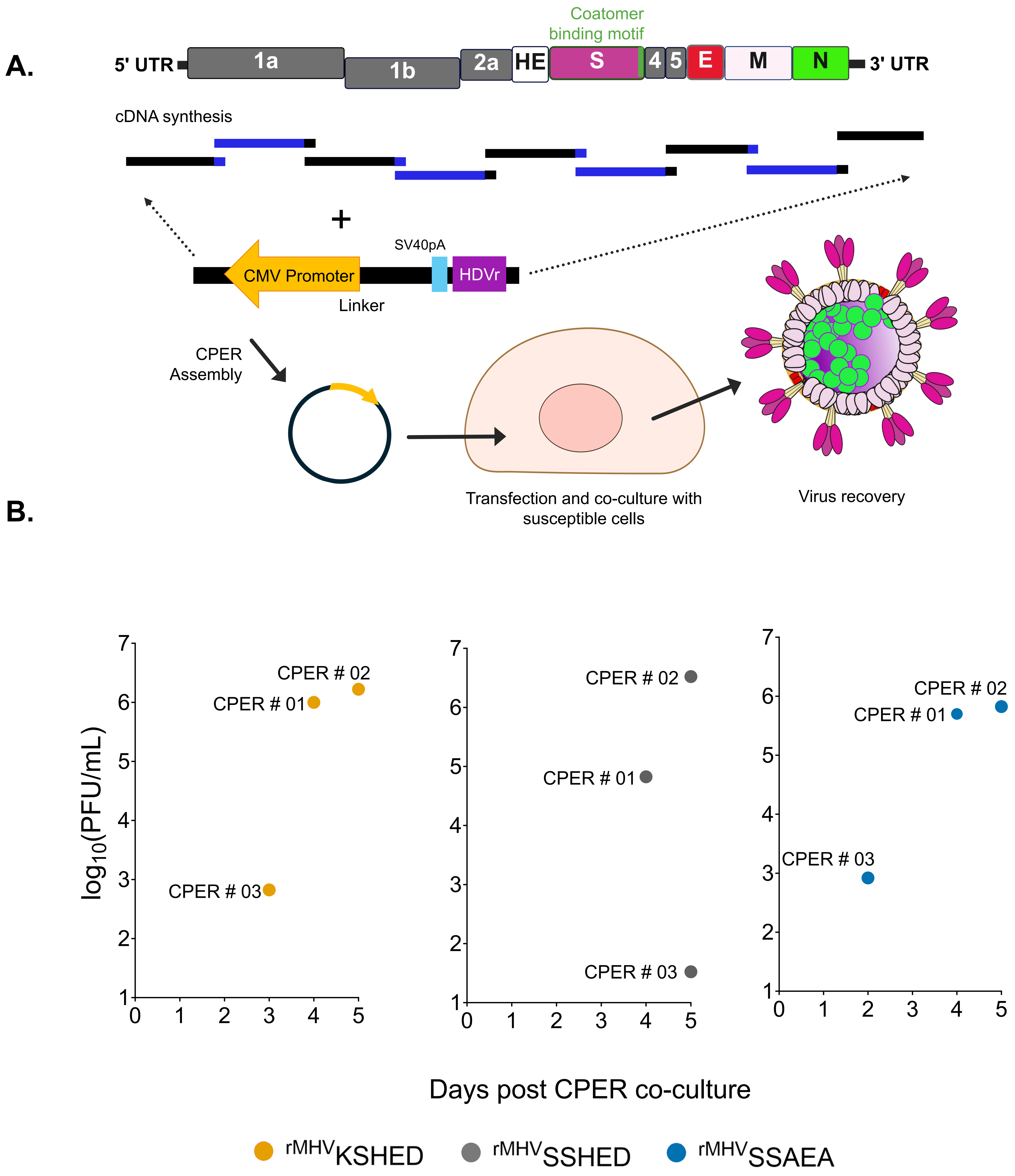


**SI Fig. S3 | S-coatomer affinity governs MHV infectivity and genetic adaptation. (A)** Schematic depicting MHV genome and rMHV generation using Circular Polymerase Extension Reaction (CPER). **(B)** Titer of recovered virus after independent CPER experiments, at different time points post co-culture (n=3). KSHED viruses recovered from CPER#1 and #2 contain second site suppressor mutations.
