## Supplementary material for "An unconventional HxD motif orchestrates coatomer-dependent coronavirus morphogenesis": SI Fig. S4

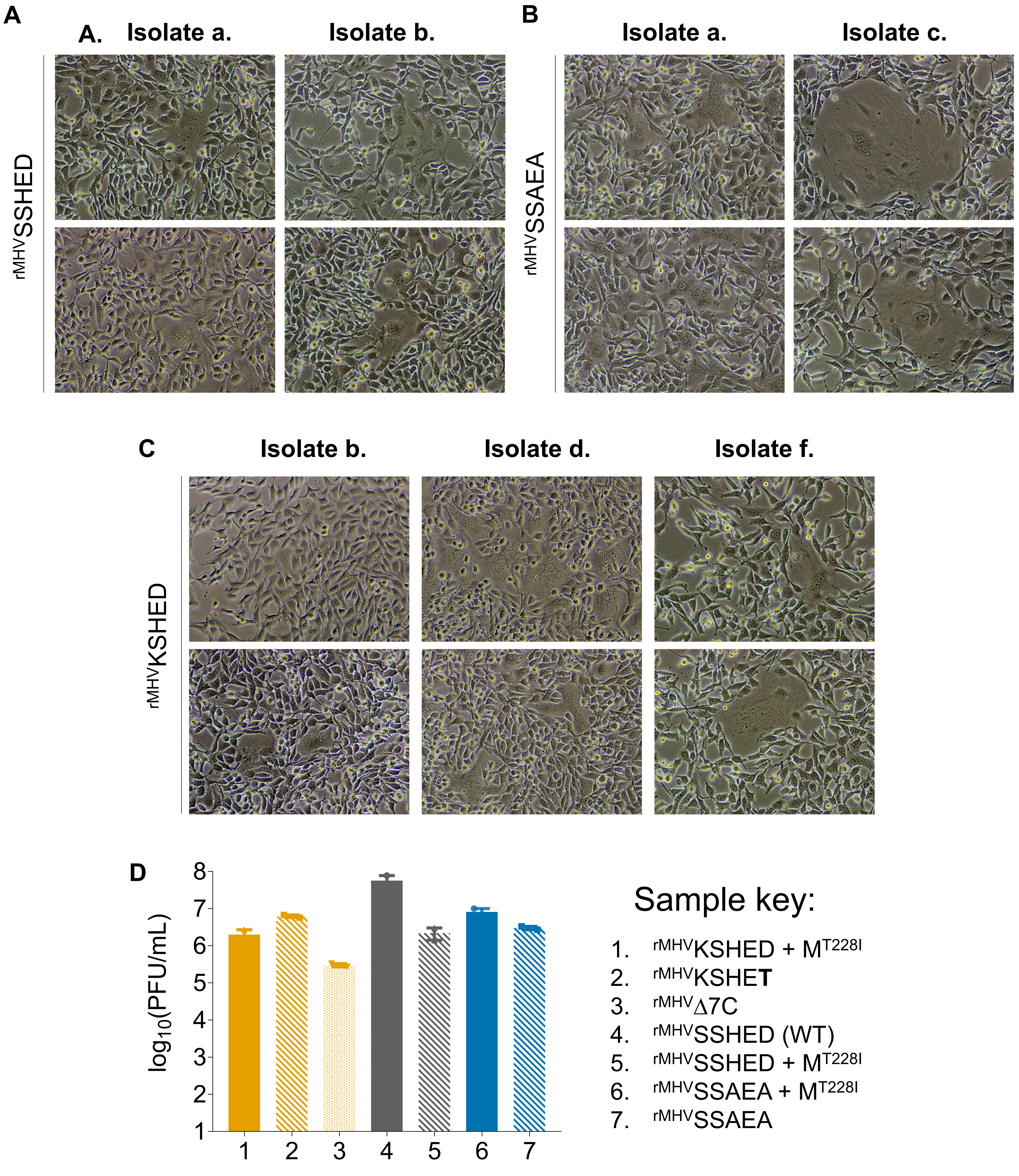


**SI Fig. S4 | Coatomer association controls plasma membrane S distribution and syncytia formation.** Phase contrast microscopy images documenting syncytia at 12 hours post infection across recombinant MHV infected 17CL1 cells. **(A)** SSHED, **(B)** SSAEA , and **(C)** KSHED virus isolates from SI table 04. Two representative regions-of-interest (ROIs) documented per rMHV. **(D)** Output titers following equal input MOI infection across recombinant MHVs from SI table 04.
