## Supplementary material for "An unconventional HxD motif orchestrates coatomer-dependent coronavirus morphogenesis": SI Table T1

**SI Table T1 | Sequence relationships between coatomer WD40 domains^1,2,3^**

|  | Sp_αWD40 | Mm_αWD40 | Hs_αWD40 | Sc_β΄WD40 | Mm_β2WD40 | Hs_β2WD40 |
| --- | --- | --- | --- | --- | --- | --- |
| Sp_αWD40 |  | 71.9/83.2 | 71.6/83.2 | 31.2/50.0 | 31.9/49.6 | 31.9/49.6 |
| Mm_αWD40 |  |  | 99.7/100 | 32.1/49.7 | 34.5/50.0 | 34.5/50.0 |
| Hs_αWD40 |  |  |  | 32.1/49.7 | 34.5/50.0 | 34.5/50.0 |
| Sc_β΄WD40 |  |  |  |  | 58.4/76.2 | 58.4/76.2 |
| Mm_ β2WD40 |  |  |  |  |  | 100.0/100.0 |

^1^Percent identity/similarity

^2^Hs, *Homo sapiens*; Mm, *Mus musculus*; Sc, *Saccharomyces cerevisiae*; Sp, *Schizosaccharomyces pombe*

^3^Sequences used

>Q96WV5_Sp_aWD40

MEMLTKFESRSSRAKGVAFHPTQPWILTSLHNGRIQLWDYRMGTLLDRFDGHDGPVRGIAFHPTQPLFVSGGDDYKVNVWNYKSRKLLFSLCGHMDYVRVCTFHHEYPWILSCSDDQTIRIWNWQSRNCIAILTGHSHYVMCAAFHPSEDLIVSASLDQTVRVWDISGLRMKNAAPVSMSLEDQLAQAHNSISNDLFGSTDAIVKFVLEGHDRGVNWCAFHPTLPLILSAGDDRLVKLWRMTASKAWEVDTCRGHFNNVSCCLFHPHQELILSASEDKTIRVWDLNRRTAVQTFRRDNDRFWFITVHPKLNLFAAAHDSGVMVFKLE

>P53621_Hs_aWD40

MLTKFETKSARVKGLSFHPKRPWILTSLHNGVIQLWDYRMCTLIDKFDEHDGPVRGIDFHKQQPLFVSGGDDYKIKVWNYKLRRCLFTLLGHLDYIRTTFFHHEYPWILSASDDQTIRVWNWQSRTCVCVLTGHNHYVMCAQFHPTEDLVVSASLDQTVRVWDISGLRKKNLSPGAVESDVRGITGVDLFGTTDAVVKHVLEGHDRGVNWAAFHPTMPLIVSGADDRQVKIWRMNESKAWEVDTCRGHYNNVSCAVFHPRQELILSNSEDKSIRVWDMSKRTGVQTFRRDHDRFWVLAAHPNLNLFAAGHDGGMIVFKLE

>O55029_Mm_b2WD40

MPLRLDIKRKLTARSDRVKSVDLHPTEPWMLASLYNGSVCVWNHETQTLVKTFEVCDLPVRAAKFVARKNWVVTGADDMQIRVFNYNTLERVHMFEAHSDYIRCIAVHPTQPFILTSSDDMLIKLWDWDKKWSCSQVFEGHTHYVMQIVINPKDNNQFASASLDRTIKVWQLGSSSPNFTLEGHEKGVNCIDYYSGGDKPYLISGADDRLVKIWDYQNKTCVQTLEGHAQNVSCASFHPELPIIITGSEDGTVRIWHSSTYRLESTLNYGMERVWCVASLRGSNNVALGYDEGSIIVKLG

>P41811_Sc_b’WD40

MKLDIKKTFSNRSDRVKGIDFHPTEPWVLTTLYSGRVELWNYETQVEVRSIQVTETPVRAGKFIARKNWIIVGSDDFRIRVFNYNTGEKVVDFEAHPDYIRSIAVHPTKPYVLSGSDDLTVKLWNWENNWALEQTFEGHEHFVMCVAFNPKDPSTFASGCLDRTVKVWSLGQSTPNFTLTTGQERGVNYVDYYPLPDKPYMITASDDLTIKIWDYQTKSCVATLEGHMSNVSFAVFHPTLPIIISGSEDGTLKIWNSSTYKVEKTLNVGLERSWCIATHPTGRKNYIASGFDNGFTVLSLG

>Q8CIE6_Mm_aWD40

MLTKFETKSARVKGLSFHPKRPWILTSLHNGVIQLWDYRMCTLIDKFDEHDGPVRGIDFHKQQPLFVSGGDDYKIKVWNYKLRRCLFTLLGHLDYIRTTFFHHEYPWILSASDDQTIRVWNWQSRTCVCVLTGHNHYVMCAQFHPSEDLVVSASLDQTVRVWDISGLRKKNLSPGAVESDVRGITGVDLFGTTDAVVKHVLEGHDRGVNWAAFHPTMPLIVSGADDRQVKIWRMNESKAWEVDTCRGHYNNVSCAVFHPRQELILSNSEDKSIRVWDMSKRTGVQTFRRDHDRFWVLAAHPNLNLFAAGHDGGMIVFKLE

>P35606_Hs_b2WD40

MPLRLDIKRKLTARSDRVKSVDLHPTEPWMLASLYNGSVCVWNHETQTLVKTFEVCDLPVRAAKFVARKNWVVTGADDMQIRVFNYNTLERVHMFEAHSDYIRCIAVHPTQPFILTSSDDMLIKLWDWDKKWSCSQVFEGHTHYVMQIVINPKDNNQFASASLDRTIKVWQLGSSSPNFTLEGHEKGVNCIDYYSGGDKPYLISGADDRLVKIWDYQNKTCVQTLEGHAQNVSCASFHPELPIIITGSEDGTVRIWHSSTYRLESTLNYGMERVWCVASLRGSNNVALGYDEGSIIVKLG
