## Supplementary material for "An unconventional HxD motif orchestrates coatomer-dependent coronavirus morphogenesis": SI Table T2

**SI Table T2 | Peptides used in this investigation^1,2^**

| **Name** | **Sequence** | **Application** |
| --- | --- | --- |
| SSHED | CDEYGGHQDSIVIHNISSHED | BLI assays |
| SSAEA | CDEYGGHQDSIVIHNISSAEA |  |
| KSHED | CDEYGGHQDSIVIHNIKSHED |  |
| KSSED | CDEYGGHQDSIVIHNIKSSED |  |
| Heptapeptide | NISSHED | Co-crystallization |

^1^BLI peptides have an N-terminal biotin group for immobilization on SA biosensors

^2^Mutations relative to wild-type SSHED 21-mer peptide are highlighted in gray
