## Supplementary material for "An unconventional HxD motif orchestrates coatomer-dependent coronavirus morphogenesis": SI Table T3

**SI Table T3 | BLI assays of MHV S tail peptides with coatomer WD40 domains**

| **Immobilized 21mer peptide** | **WD40 domain** | **KD (µM)** |
| --- | --- | --- |
| SSHED | β2 | No binding |
| SSHED | α-Fab17 | 0.50 ± 0.07^1^ |
| SSAEA | α-Fab17 | No binding |
| SSHED | α(H31A)-Fab17 | No binding |

^1^Average of three independent measurements- 0.69 ± 0.10, 0.42 ± 0.04, 0.40 ± 0.04 µM
