## Supplementary material for "An unconventional HxD motif orchestrates coatomer-dependent coronavirus morphogenesis": SI Table T4

**SI Table T4 | Crystallographic data collection and refinement statistics**

| **Protein** | **αWD40** | **αWD40 (H31A)** | **β2WD40** |
| --- | --- | --- | --- |
| Peptide | NISSHED | **--** | NISSHED |
| PDB ID | 9DZX | 9YIZ | 9YIY |
| **Data collection^1^** | | | |
| Space group | P 21 21 21 | P 1 21 1 | P 1 21 1 |
| Unit cell (a, b, c; Å) | 38.1, 75.1, 129.3 | 36.6, 172.1, 70.9 | 57.3, 93.1, 57.9 |
| Unit cell (α, β, γ; °) | 90, 90, 90 | 90, 98.7, 90 | 90, 100.7, 90 |
| Resolution (Å) | 32.8-1.66 (1.72-1.66) | 29.5-1.72 (1.78-1.72) | 28.86-1.36 (1.41-1.36) |
| R_merge_ | 0.037 (0.77) | 0.052 (0.367) | 0.079 (0.630) |
| I/σ(I) | 9.99 (0.77) | 9 (2.08) | 10.29 (1.96) |
| CC1/2 | 0.99 (0.50) | 0.99 (0.68) | 0.99 (0.675) |
| Total number of reflections | 88647 (8204) | 180970 (18265) | 513981 (44443) |
| Number of unique reflections | 44353 (4121) | 91526 (9166) | 127400 (12561) |
| Completeness (%) | 99.19 (93.68) | 99.59 (99.93) | 99.69 (98.28) |
| Wilson B factor (Å^2^) | 22.51 | 16.90 | 13.13 |
| **Refinement** | | | |
| Resolution used in refinement (Å) | 32.8-1.66 (1.72-1.66) | 29.5-1.72 (1.78-1.72) | 28.86-1.36 (1.41-1.36) |
| Number of WD40 chains in ASU | 1 | 3 | 2 |
| Number of peptide chains in ASU | 1 | -- | 1 |
| Number of reflections used in refinement | 44348 (4120) | 91472 (9166) | 127327 (12534) |
| R_work_ | 0.17 (0.37) | 0.16 (0.23) | 0.15 (0.25) |
| R_free_^2^ | 0.20 (0.39) | 0.20 (0.29) | 0.17 (0.29) |
| Amino acid residues in final model | 308 | 918 | 599 |
| Water molecules | 227 | 929 | 622 |
| All atom clash-score | 4.19 | 1.67 | 1.79 |
| Ramachandran angles (favored, %) | 95.03 | 95.47 | 95.95 |
| Ramachandran angles (allowed, %) | 4.97 | 4.42 | 4.05 |
| Ramachandran angles (outliers, %) | 0.00 | 0.11 | 0.00 |
| Rotamers outliers | 0.37 | 0.64 | 0.00 |
| RMS bond length (Å) | 0.007 | 0.010 | 0.008 |
| RMS bond angle (°) | 0.95 | 1.09 | 1.04 |

^1^Data collected at NSLSII 17-ID-1 AMX beamline at an X-ray wavelength of 0.92Å. ^2^ 5% of reflections assigned for R_free_ calculations.
