## Supplementary material for "An unconventional HxD motif orchestrates coatomer-dependent coronavirus morphogenesis": SI Table T5

**SI Table T5 | Genetic adaptation by rMHVs with altered S-coatomer affinity.**

| ENGINEERED | RECOVERED | | | | |
|---|---|---|---|---|---|
| S: C-termini | | S: C termini | NS2a | E | M |
| $^{rMHV}$SSHED (intermediate coatomer binding) | a. | ...$^{1311}$QDSIVIHNI**SSHED**$^{1324}$* | nd | nd | nd |
| | b. | | nd | F18V | T228I |
| $^{rMHV}$SSAEA (no coatomer binding) | a. | ...$^{1311}$QDSIVIHNI**SSAEA**$^{1324}$* | nd | nd | nd |
| | b. | | nd | nd | F166S T228I |
| | c. | | nd | L6P | T228I |
| $^{rMHV}$KSHED (strong binding) | a. | ...$^{1311}$QDSIVIHNI**KSHED**$^{1324}$* | Y238H | nd | nd |
| | b. | ...$^{1311}$QDSIVIHNI**KSHED**$^{1324}$* | | | T228I |
| | c. | ...$^{1311}$QDSIVIHNI**KS**Q**ED**$^{1324}$* | | nd | nd |
| | d. | ...$^{1311}$QDSIVIHNI**KSHE**T$^{1324}$* | | V30A | nd |
| | e. | ...$^{1311}$QDSIVIHNIKSH$^{1322}$* | nd | nd | nd |
| | f. | ...$^{1311}$QYCDT$^{1317}$* | Δ42-260 | nd | nd |

Sequence analysis of recovered rMHVs following CPER assembled MHV cDNA transfection. *nd = none detected.
