## Supplementary material for "An unconventional HxD motif orchestrates coatomer-dependent coronavirus morphogenesis": SI Table T6

**SI Table T6 | Coatomer-dependent modification of S content in MHV particles imaged by negative staining TEM^1^**

| **Number of S molecules** | **SSHED** | | **SSAEA** | |
| --- | --- | --- | --- | --- |
|  | **Number of particles** | **Percentage** | **Number of particles** | **Percentage** |
| 0 | 12 | 3.0 | 156 | 35.6 |
| 1 | 7 | 1.8 | 53 | 12.1 |
| 2 | 6 | 1.5 | 35 | 8 |
| 3 | 7 | 1.8 | 63 | 14.4 |
| 4 | 13 | 3.3 | 31 | 7.1 |
| 5 | 25 | 6.3 | 23 | 5.3 |
| 6 | 45 | 11.4 | 28 | 6.4 |
| 7 | 39 | 9.9 | 15 | 3.4 |
| 8 | 30 | 7.6 | 7 | 1.6 |
| 9 | 36 | 9.1 | 9 | 2.1 |
| 10 | 31 | 7.8 | 4 | 0.9 |
| 11 | 34 | 8.6 | 3 | 0.7 |
| 12 | 31 | 7.8 | 5 | 1.1 |
| 13 | 19 | 4.8 | 3 | 0.7 |
| 14 | 23 | 5.8 | 2 | 0.5 |
| 15 | 9 | 2.3 | 1 | 0.2 |
| 16 | 11 | 2.8 | 0 | 0 |
| 17 | 6 | 1.5 | 0 | 0 |
| 18 | 5 | 1.3 | 0 | 0 |
| 19 | 4 | 1.0 | 0 | 0 |
| 20 | 0 | 0.0 | 0 | 0 |
| 21 | 0 | 0.0 | 0 | 0 |
| 22 | 2 | 0.5 | 0 | 0 |
| Total | 395 | 100 | 438 | 100 |

^1^Heat map- red to blue from most to fewest particles.
